# Nonlinear Dispersive Cell Model for Microdosimetry of Nanosecond Pulsed Electric Fields

**DOI:** 10.1101/660134

**Authors:** Fei Guo, Lin Zhang, Xin Liu

## Abstract

For nanosecond pulsed electric fields (nsPEFs) based application, the underlying transmembrane potential (TMP) distribution of the plasma membrane is influenced by electroporation (EP) of the plasma membrane and dispersion (DP) of all cell compartments and is important for predicting the bioelectric effects. In this study, we analysed temporal and spatial distribution of TMP induced by nsPEFs of various durations (3 ns, 5 ns unipolar, 5 ns bipolar, and 10 ns) with the consideration of both DP and EP. Based on the double-shelled dielectric spherical cell model, we used second-order Debye equation to characterize the dielectric relaxation of plasma membrane and nuclear membrane in the frequency domain and transformed the Debye equation into the time domain with the introduction of polarization vector, then we obtained the time course of TMP by solving the combination of Laplace equation and time-domain Debye equation. Next, we used the asymptotic version of the smoluchowski equation to characterize electroporation of plasma membrane and added it to our model to achieve the temporal and spatial distribution of TMP and pore density. Much faster and more pronounced increased in TMP can be found with the consideration of dielectric relaxation of plasma membrane and nuclear membrane, and much larger electroporated area of at least half of the plasma membrane was obtained with the consideration of both DP and EP. Through the simulation it is clearer to understand the relationship.

## Introduction

Transmembrane potential (TMP) appears on the plasma membrane when a biological cell is exposed to external electric fields. If the external filed intensity is strong, TMP will exceed the physiological range of the potential on the plasma membrane (0.4-1V). In this situation, micro-pores occur on the membrane, and this phenomenon is called electroporation (EP) [1–2]. EP has become a common method for gene transfection, drug delivery, and been studying for cancer treatment [3–6].

Typically, EP uses pulse electric fields with the field intensity of several kV/cm and the duration in the level of several hundred of microseconds to several milliseconds [1–3]. Recently, electric pulses with the field intensity of several tens of kV/cm and duration in the level of nanoseconds have been regarded as a drug free, non-thermal way to address cancer diseases [7–10]. Both model evidences and experimental results indicate that nsPEFs induce structural and functional changes of intracellular organelles, which is different from traditional electroporation [11–15]. Compare with conventional EP, much more numerous, but smaller-sized pores are created in almost all regions of the plasma membrane with the application of intense nsPEFs [16], which induced a significant increase in conductivity of the plasma membrane during and after nsPEFs exposure [17–18], the appearance of massive micro-pore and secondary effects are closely related to the distribution of TMP of plasma membrane, therefore, accurately calculation of TMP of plasma membrane plays a critical role in predicting the desired biological effects [19–20].

However, it is difficult to directly observe the changes of TMP on the plasma membrane in real-time during nsPEFs exposure. The study of the relationship between nsPEFs and TMP commonly relies on theoretical analysis. In previous theoretical studies, two effects that were always ignored can greatly affect the temporal and spatial distribution of TMP when application of nsPEFs to biological cells: 1) dielectric dispersion (DP), conductivity and permittivity of each component of a biological cell is frequency-dependent, in consequences TMP of a biological cell depends on frequency spectrum of the applied nsPEFs [20–24]; 2) electroporation, micro-pores occurred on the plasma membrane greatly increases its conductivity, then the distribution of TMP will be changed [25–28]. Smoluchowski equation was used to investigate the creation and development of micro-pores on the plasma membrane in previous studies when studying the effect of electroporation on TMP of a biological cell [28]. The effects of dielectric dispersion of cell components on the TMP of plasma membrane were investigated both in the time and frequency domain [20, 22–25]. To the best of our knowledge, few studies have investigated the effects of both DP and EP on the temporal and spatial distribution of TMP of plasma membrane. A quasi-static solution based on Laplace equation was adapted to nsPEFs and the electric solution then was coupled with an asymptotic electroporation model to investigate the effects of both EP and DP in [22], the calculation involved a two-step process and cannot obtain the effects of both EP and DP on TMP simultaneously. Joshi and colleagues presented the time-dependent transmembrane potential at the outer cell membrane with the consideration of both EP and DP, based on the numerical distribution circuit approach [24]. In [25], Salimi and colleagues investigated membrane dielectric dispersion in nanosecond pulsed electroporation of biological cells, based on the single-shelled cell model.

An improved method based on [25], both DP and EP can be easily investigated simultaneously with the introduction of polarization vector, which is very convenient for us to investigate the temporal and spatial distribution of TMP based on the double-shelled cell model, is presented in this study.

## Materials and methods

In Section II-A, the dielectric double-shelled cell model and cell geometric characteristics are given, followed by the description of the Debye dispersive model in Section II-B. The asymptotic model of electroporation and the pulse characteristics are briefly outlined in Sections II-C and II-D, respectively. Finally, the model setting and calculations of the induced TMP and pore density are explained in Section II-E.

### A. Dielectric double-shelled cell model

A sphere contained a smaller sphere inside, was developed as the dielectric double-shelled cell model and was adopted in our study, as shown in Fig 1. The large and small spheres were all shielded by thin layers (represents the plasma membrane or nuclear membrane). Each component of this model was assumed to be isotropy. To analyse the evolution of pore density and transmembrane potential on the surface of the cell membrane, seven sampling points (A_1_-A_7_) were selected, and the angle between every next two points was 15°. The parameters of this model are detailed in Table 1.

**Fig 1.**
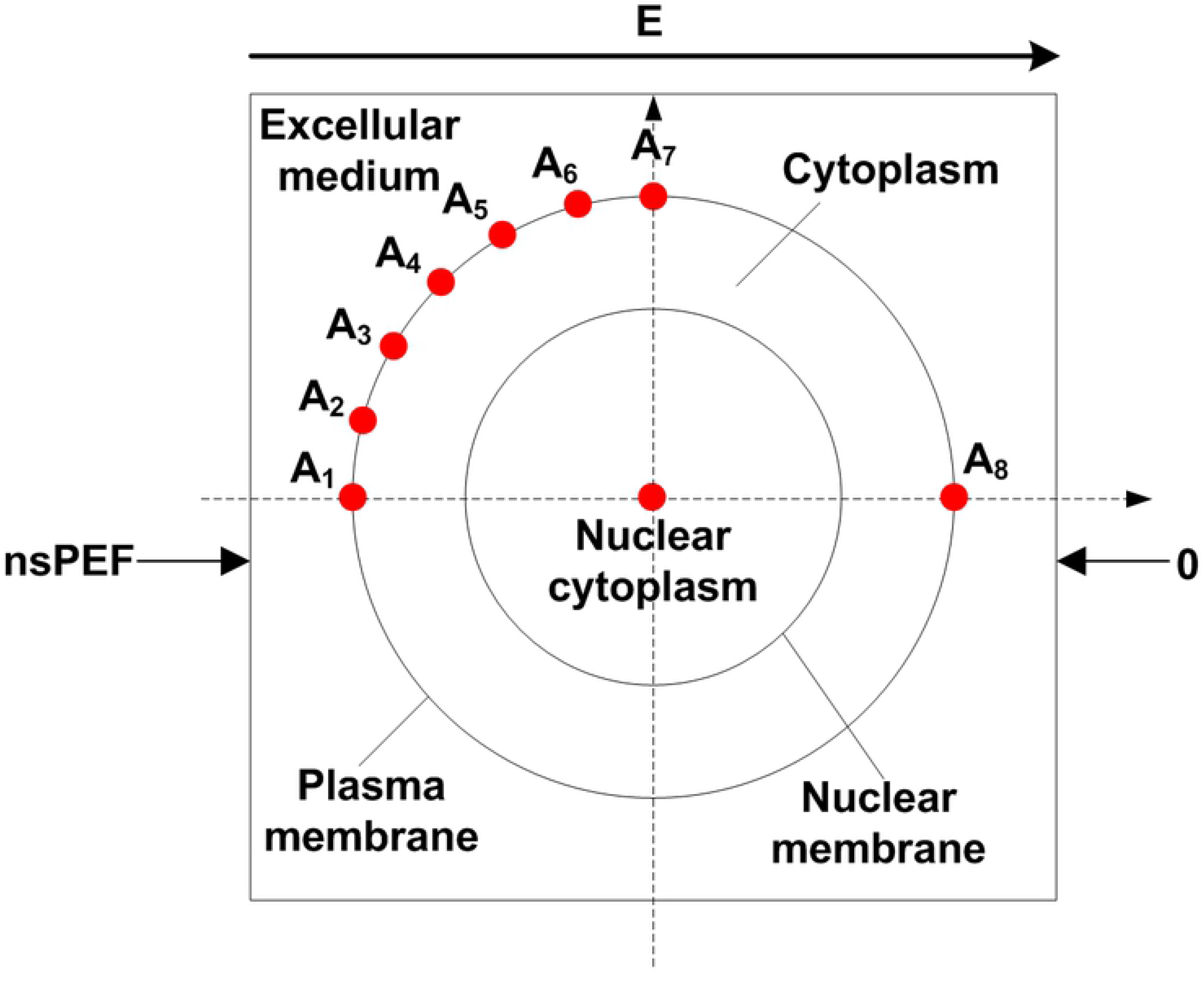
Dielectric double-shelled cell model.

**Table 1.**
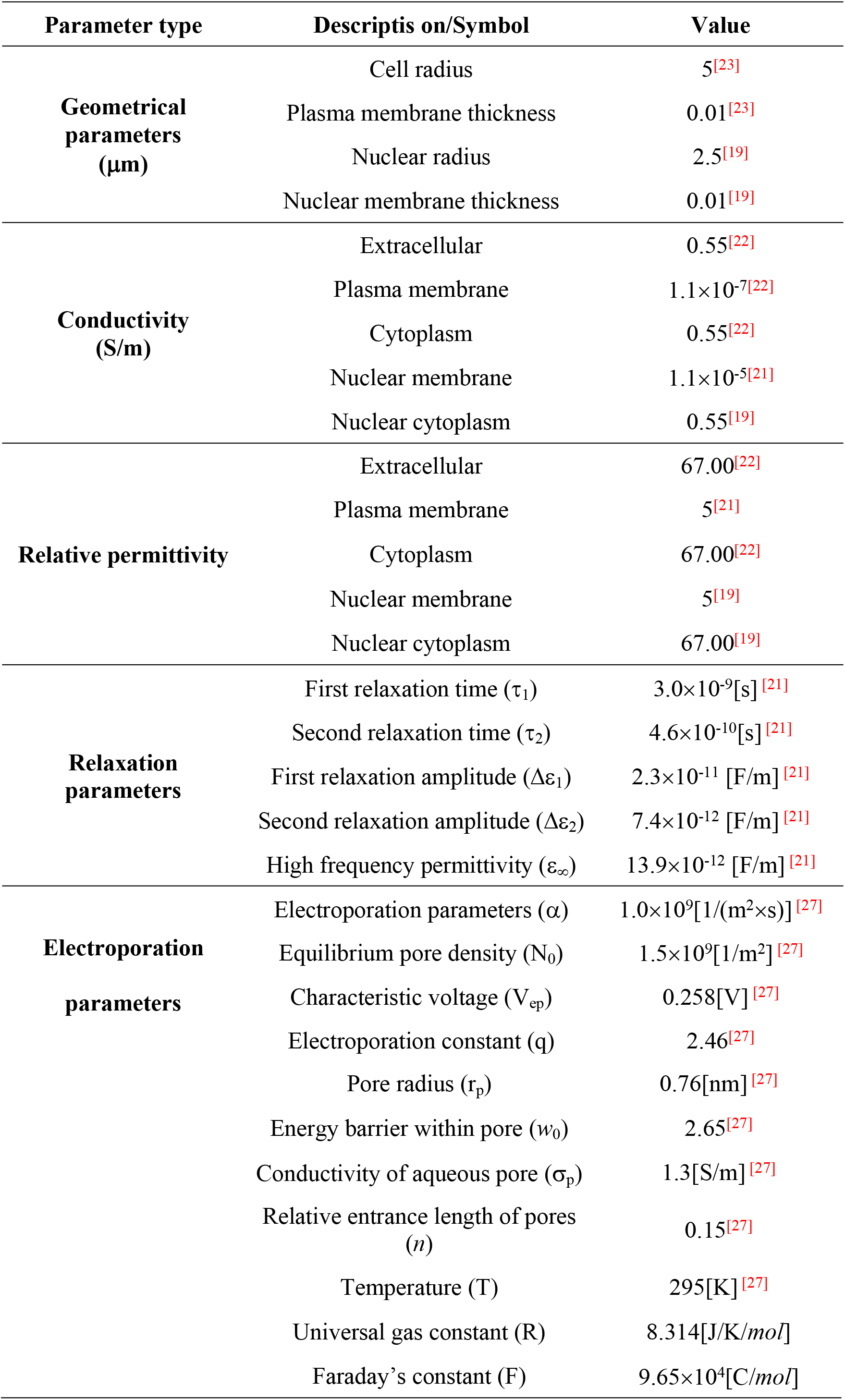
Cell paramters used in our study

| Parameter type | Descriptis on/Symbol | Value |
| --- | --- | --- |
| **Geometrical parameters**<br>($\mu\text{m}$) | Cell radius | 5<sup>[23]</sup> |
|  | Plasma membrane thickness | 0.01<sup>[23]</sup> |
|  | Nuclear radius | 2.5<sup>[19]</sup> |
|  | Nuclear membrane thickness | 0.01<sup>[19]</sup> |
| **Conductivity**<br>(S/m) | Extracellular | 0.55<sup>[22]</sup> |
| | Plasma membrane | $1.1 \times 10^{-7}$<sup>[22]</sup> |
|  | Cytoplasm | 0.55<sup>[22]</sup> |
| | Nuclear membrane | $1.1 \times 10^{-5}$<sup>[21]</sup> |
|  | Nuclear cytoplasm | 0.55<sup>[19]</sup> |
| **Relative permittivity** | Extracellular | 67.00<sup>[22]</sup> |
|  | Plasma membrane | 5<sup>[21]</sup> |
|  | Cytoplasm | 67.00<sup>[22]</sup> |
|  | Nuclear membrane | 5<sup>[19]</sup> |
|  | Nuclear cytoplasm | 67.00<sup>[19]</sup> |
| **Relaxation parameters** | First relaxation time ($\tau_1$) | $3.0 \times 10^{-9}$[s]<sup>[21]</sup> |
| | Second relaxation time ($\tau_2$) | $4.6 \times 10^{-10}$[s]<sup>[21]</sup> |
| | First relaxation amplitude ($\Delta\epsilon_1$) | $2.3 \times 10^{-11}$ [F/m]<sup>[21]</sup> |
| | Second relaxation amplitude ($\Delta\epsilon_2$) | $7.4 \times 10^{-12}$ [F/m]<sup>[21]</sup> |
| | High frequency permittivity ($\epsilon_\infty$) | $13.9 \times 10^{-12}$ [F/m]<sup>[21]</sup> |
| **Electroporation parameters** | Electroporation parameters ($\alpha$) | $1.0 \times 10^9$[1/(m<sup>2</sup>×s)]<sup>[27]</sup> |
| | Equilibrium pore density ($N_0$) | $1.5 \times 10^9$[1/m<sup>2</sup>]<sup>[27]</sup> |
| | Characteristic voltage ($V_{ep}$) | 0.258[V]<sup>[27]</sup> |
|  | Electroporation constant (q) | 2.46<sup>[27]</sup> |

|  |  |
| --- | --- |
| Pore radius ($r_p$) | 0.76[nm] <sup>[27]</sup> |
| Energy barrier within pore ($w_0$) | 2.65 <sup>[27]</sup> |
| Conductivity of aqueous pore ($\sigma_p$) | 1.3[S/m] <sup>[27]</sup> |
| Relative entrance length of pores<br>($n$) | 0.15 <sup>[27]</sup> |
| Temperature (T) | 295[K] <sup>[27]</sup> |
| Universal gas constant (R) | 8.314[J/K/mol] |
| Faraday's constant (F) | 9.65×10<sup>4</sup>[C/mol] |

### B. Debye dispersion

The static cell model was often treated as frequency-independent, and the cellular components should be regarded as lossy dielectrics when the applied electric field with frequency higher than megahertz. Commonly, effective conductivity and effective dielectric permittivity were used to describe their changes with frequency. Second-order Debye equation, which described the complex permittivity, was used in calculation of TMP in the time domain. The equation is expressed as:

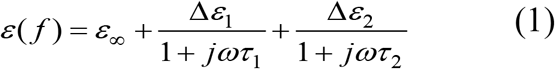

For a linear and isotropic medium the polarization vector is expressed as:

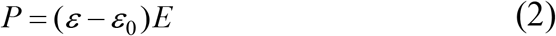

Where *ε* and *ε*_0_ are the permittivity of the medium and vacuum, respectively. Dispersion is accomplished in the time-domain by defining the polarization of the medium as a function of the electric field and its time derivatives. For a second order dispersive medium substitution of (1) into (2) and taking *jω* as the derivative with respect to time yields.

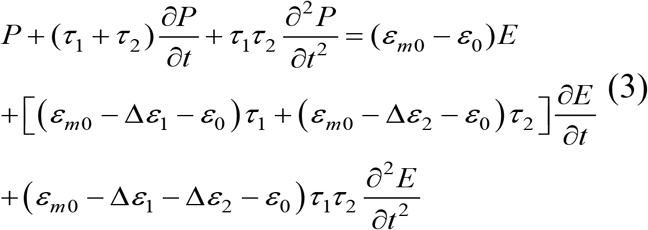

Where *ε*_m0_ is the low frequency permittivity of the membrane.

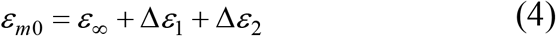

### C. Electroporation Equation

The electroporation model used here is the asymptotic version of the Smoluchoski equation, and this model are plausible for signal durations in the nanosecond time scale as noted in [29–30]. Equation 5 describes the rate of creation and destruction of hydrophilic membrane pores per local membrane area *N*(*t*) as a function of the TMP(*t*).

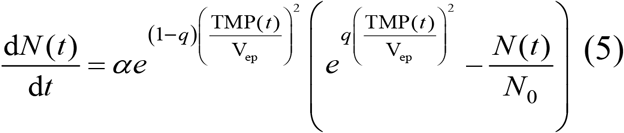

Where TMP(*t*) is the transmembrane potential of plasma membrane, the definitions and typical values of the constants in (1) - (5) are given in Table 1.

### D. Features of the nsPEFs

Trapezoidal-shaped pulses were adopted, as suggested in [21]. The pulse durations were 10 ns and 3 ns with amplitude of 10 and 18.3 kV/cm, respectively. In addition, bipolar pulse of pulse duration of 5 ns and interval of 6 ns, and unipolar pulse of pulse duration of 5 ns and interval of 6 ns were adopted, with amplitude of 10 kV/cm. All pulses have the same power density to obtain comparable results. The rise and fall times were chosen to equal to 1 ns for all pulses (Fig 2).

**Fig 2.**
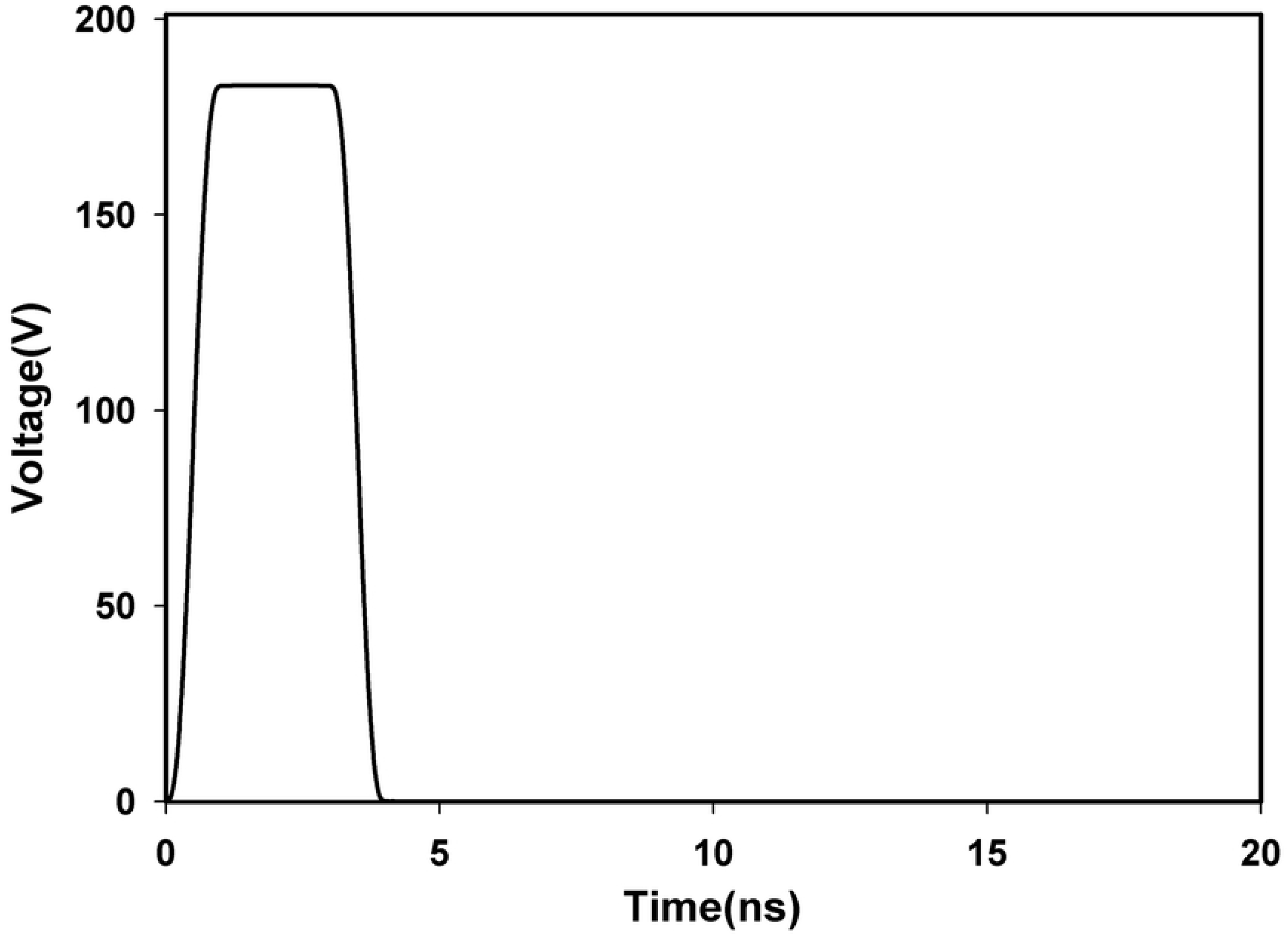
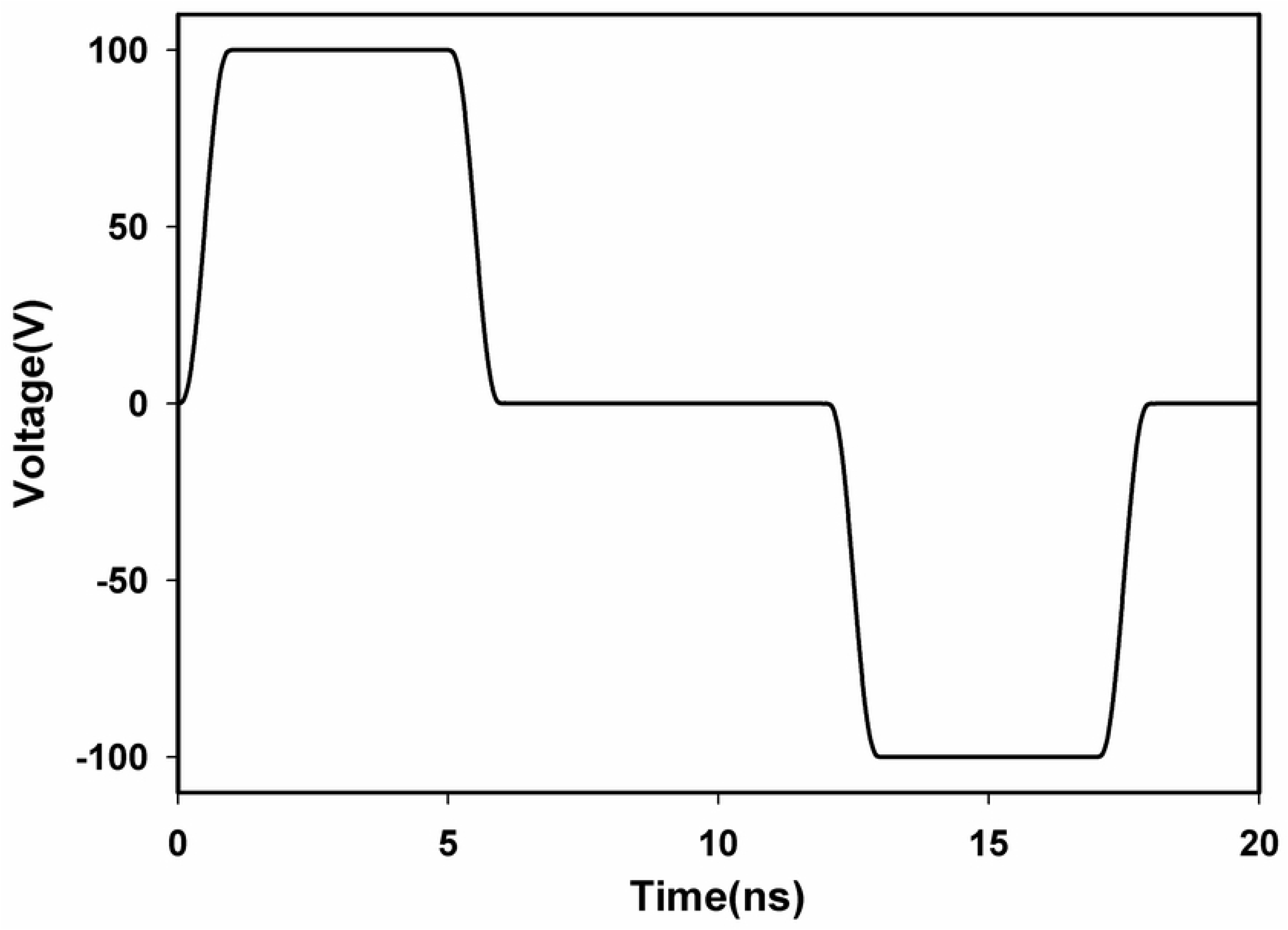
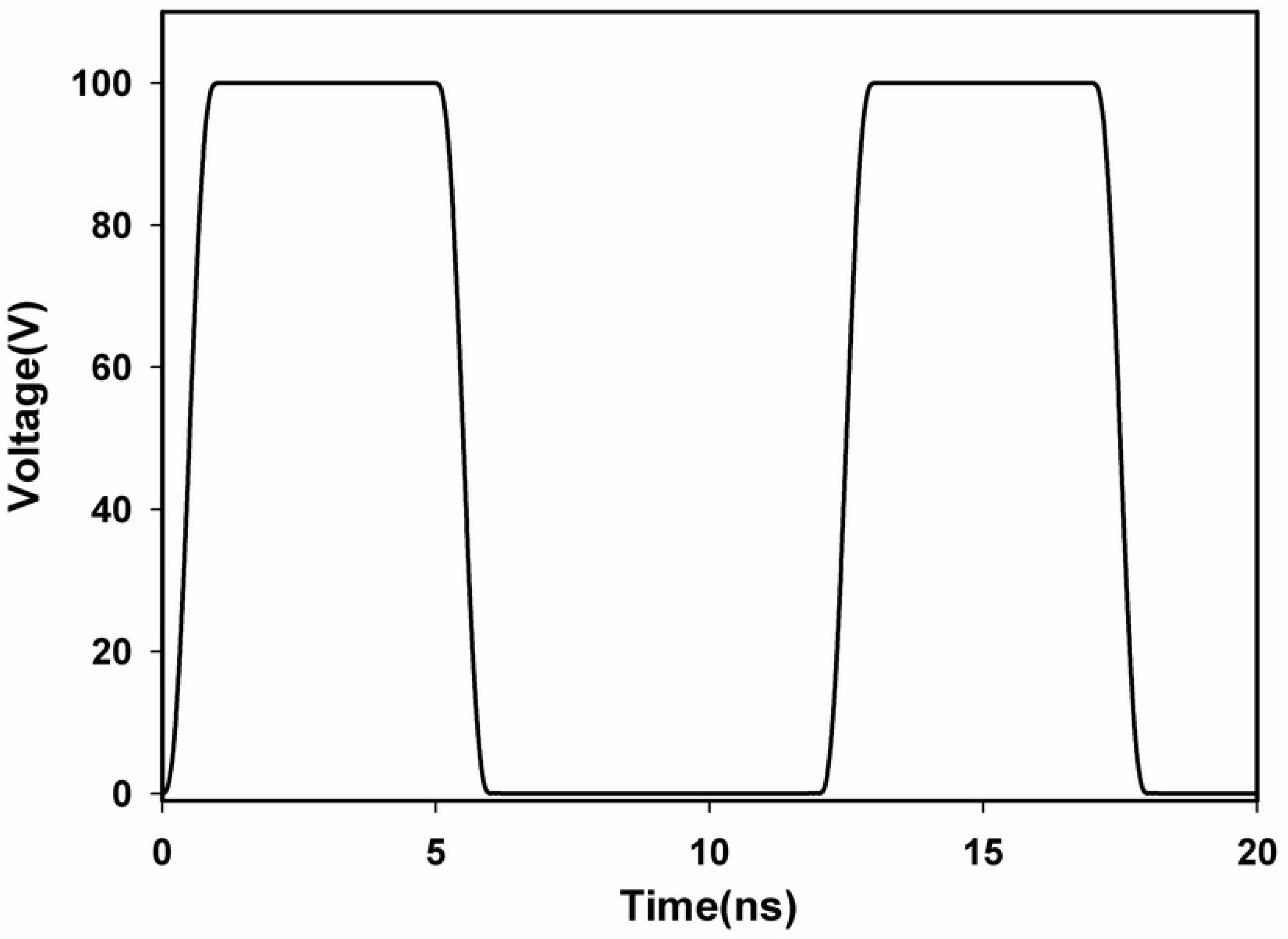
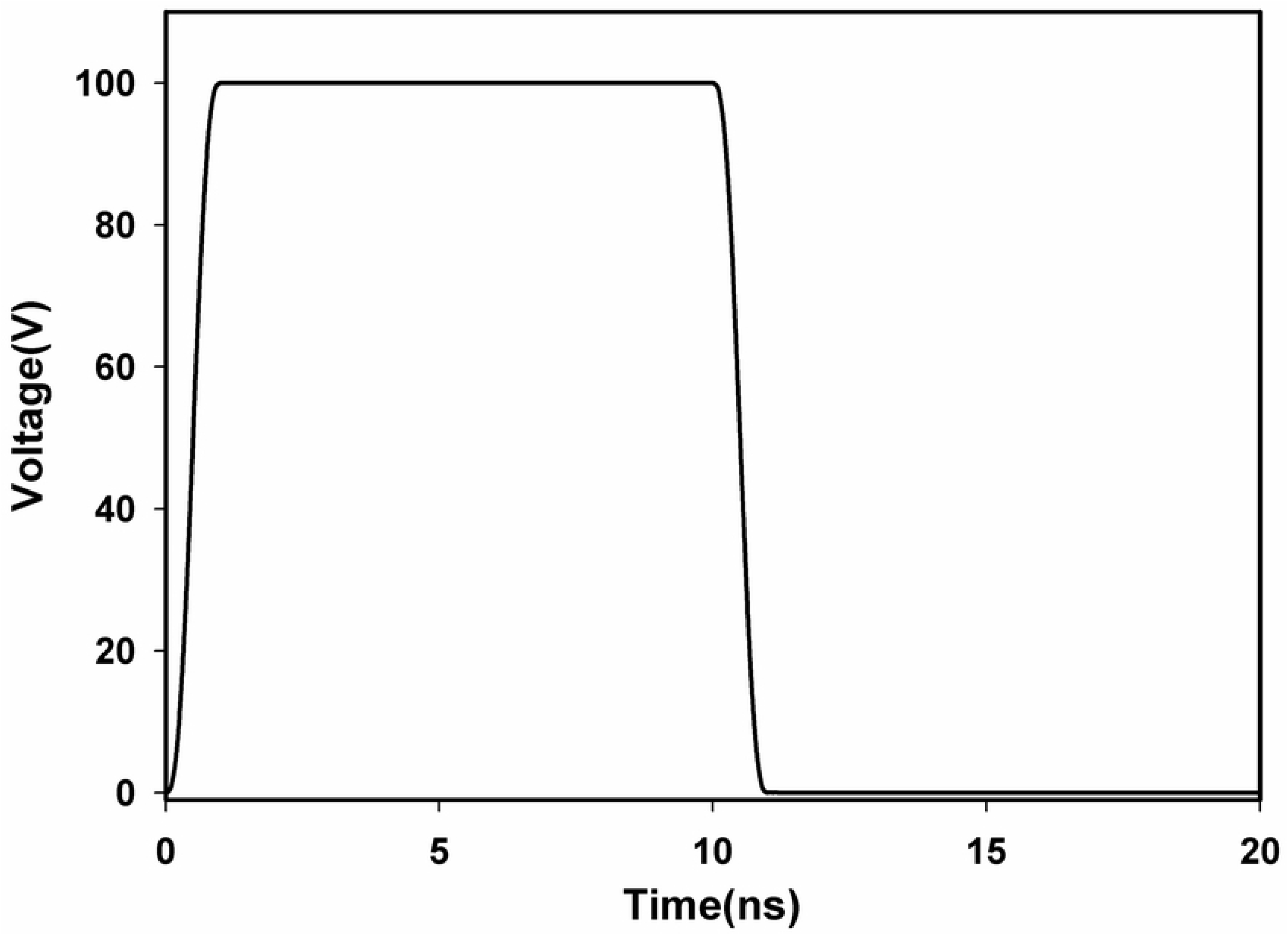
Modeled electrical pulse shapes, magnitudes, and pulse width. Single nsPEFs of duration of 3 ns and amplitude of 183 V (a), bipolar nsPEFs of pulse duration of 5 ns with time interval of 6ns and amplitude of 100V (b), unipolar nsPEFs of pulse duration of 5 ns with time interval of 6ns and amplitude of 100 V (c), single nsPEFs of duration of 10ns and amplitude of 100V (d). For 3 ns pulse, the ratio of voltage to distance is 18.3 kV/cm, and for the latter three pulses, which is 10 kV/cm, to ensure the same power density within all cases for comparison.

### E. Model settings and calculation of the induced TMP

The calculations were performed in Comsol Multiphysics 5.3a using the Electric currents, and the PDE modes-coefficient form, transient analysis mode. The opposite vertical faces of the block were modelled as electrodes, which was done by assigning electric potential to each face. The right electrode was set to electric pulse of duration of 10 and 3 ns (or 5 ns bipolar and unipolar pulses) and the left to the ground to obtain the desired electric field. The remaining faces of the block were modelled as insulating. The mesh size was refined until there was less than a 2% difference in the field results between refinements, resulting in fine mesh setting. The electric potential ϕ inside and outside the cell was then computed by solving the equation.

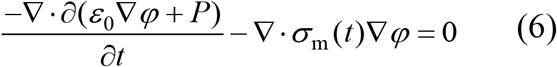

We use Electric Currents to solve the Laplace equation, the PDE modes-coefficient form to solve the pore density equation and the polarization vector equation. The Laplace equation is solved at the subdomains of extracellular medium, plasma membrane, cytoplasm, nuclear membrane and nuclear cytoplasm, the pore density equation is solved on the subdomain of plasma membrane, and the polarization vector equation is solved inside the subdomains of plasma membrane and nuclear membrane, the initial value of all the variables are set to zero at t=0 except for the resting potential is set to −70 mV and the initial density of the pores on the plasma membrane which is set to N_0_, the equilibrium pore density. Finally, the induced transmembrane potential was calculated as the difference between electric potentials on both sides of the membrane:

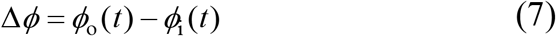

and was plotted as a function of the arc length and time.

## Results

Results are subdivided into three sections. At first, simulation verification by comparing the simulation results with analytical results is described, and then the distribution of TMP with Debye dispersive model is investigated. Finally, time evolution and spatial distribution of TMP and pore density with the consideration of both DP and EP is studied.

### A. Simulation verification

To test the accuracy of the Comsol Multiphysics code, based on a dielectric double-shelled cell model without either DP or EP, we examined the TMP of point A_1_(where TMP is maximum) with the electric field of pulse duration of 100 μs and field intensity of 1 kV/cm by comparing the analytical and simulation results. The analytical result was done by solving the first-order Schwan equation with parameters in Table 1. Fig 3 shows the time evolution of TMP of point A_1_, and the simulation result agrees very well with the analytical result, yet, the analytical result is a bit larger between 5 μs and 105 μs, which could be due to zero permittivity considered in first-order Schwan equation while our simulation did include a finite permittivity. But in general, the temporal trend of the simulation and analytical results is similar, so we think the simulation has a satisfactory accuracy. The reason why we used the cell model without either DP or EP is that the analytical result of TMP in such cell model is so complicated. Furthermore, pulsed electric field of duration of 100 μs instead of 100 ns was used because the time-domain result of TMP with the duration time of pulsed electric field less than the charging time of plasma membrane (~1 μs) is complicated.

**Fig 3.**
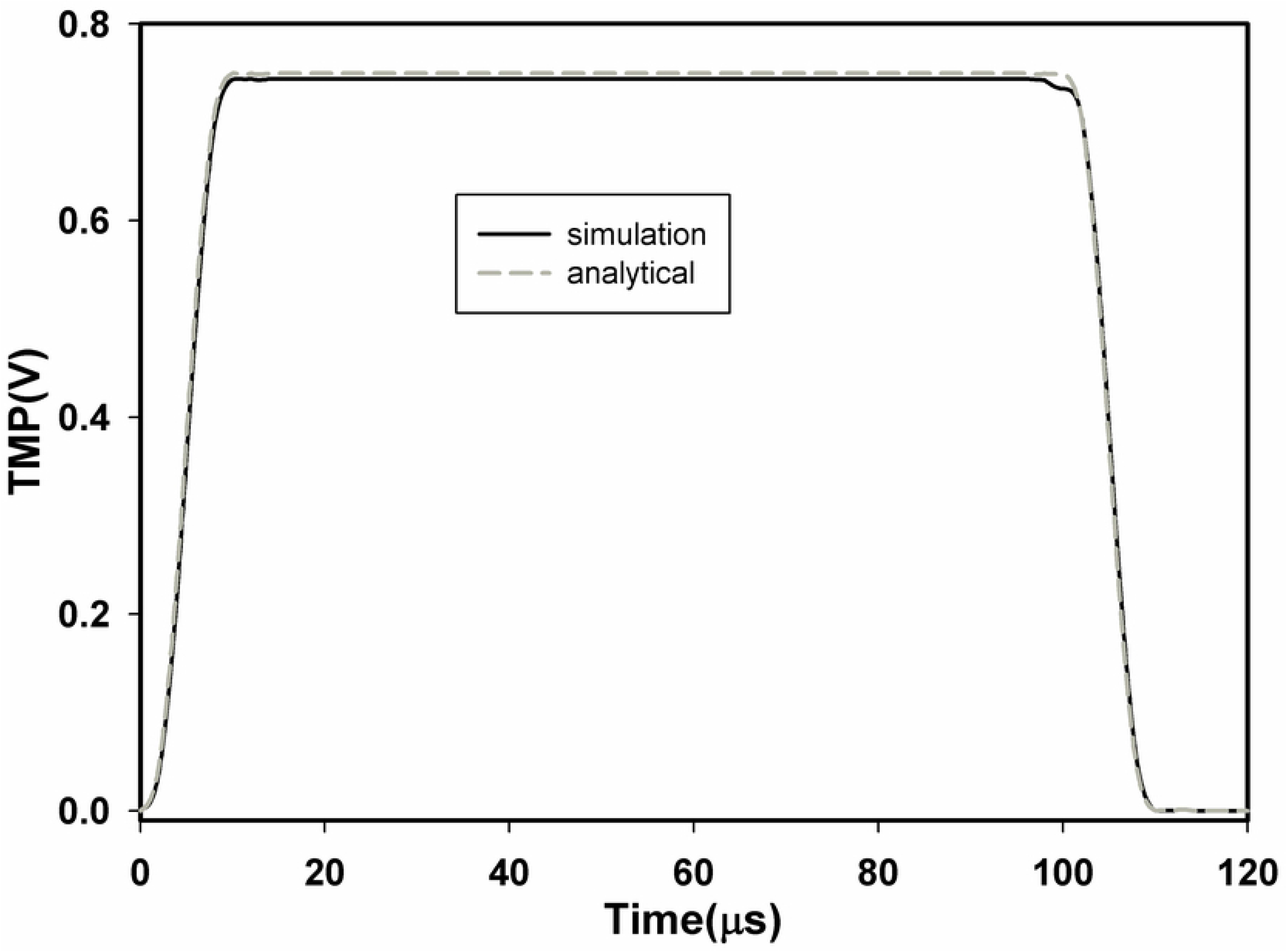
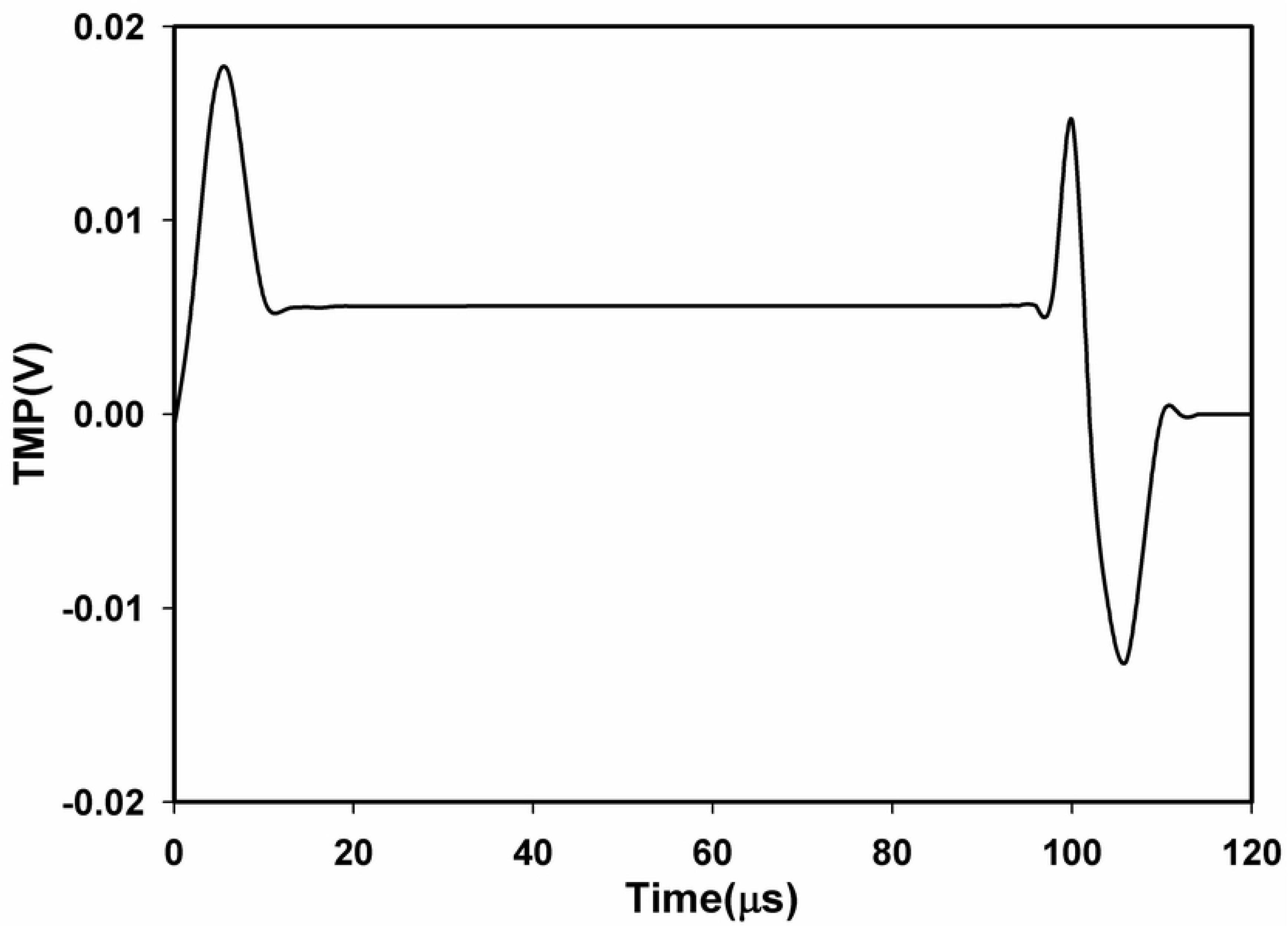
Time evolution of TMP of point A_1_. Time evolution of TMP of A_1_ with μsPEF of 100 μs and 1 kV/cm (a) and the error between analytical and simulation results, in which the analytical result obtained by solving the first-order Schwan equation (b).

### B. TMP distribution with and without dielectric relaxation

First, we investigated the TMP distribution of plasma membrane and nuclear membrane in the frequency domain when the amplitude of the electric field is 10 kV/cm in two different modes, with and without DP, and the results are shown in Fig 4a. TMP of the plasma membrane shows first order low-pass filter characteristic, while nuclear membrane shows first-order band-pass filter characteristic approximately, which agrees well with previous studies [31].

**Fig 4.**
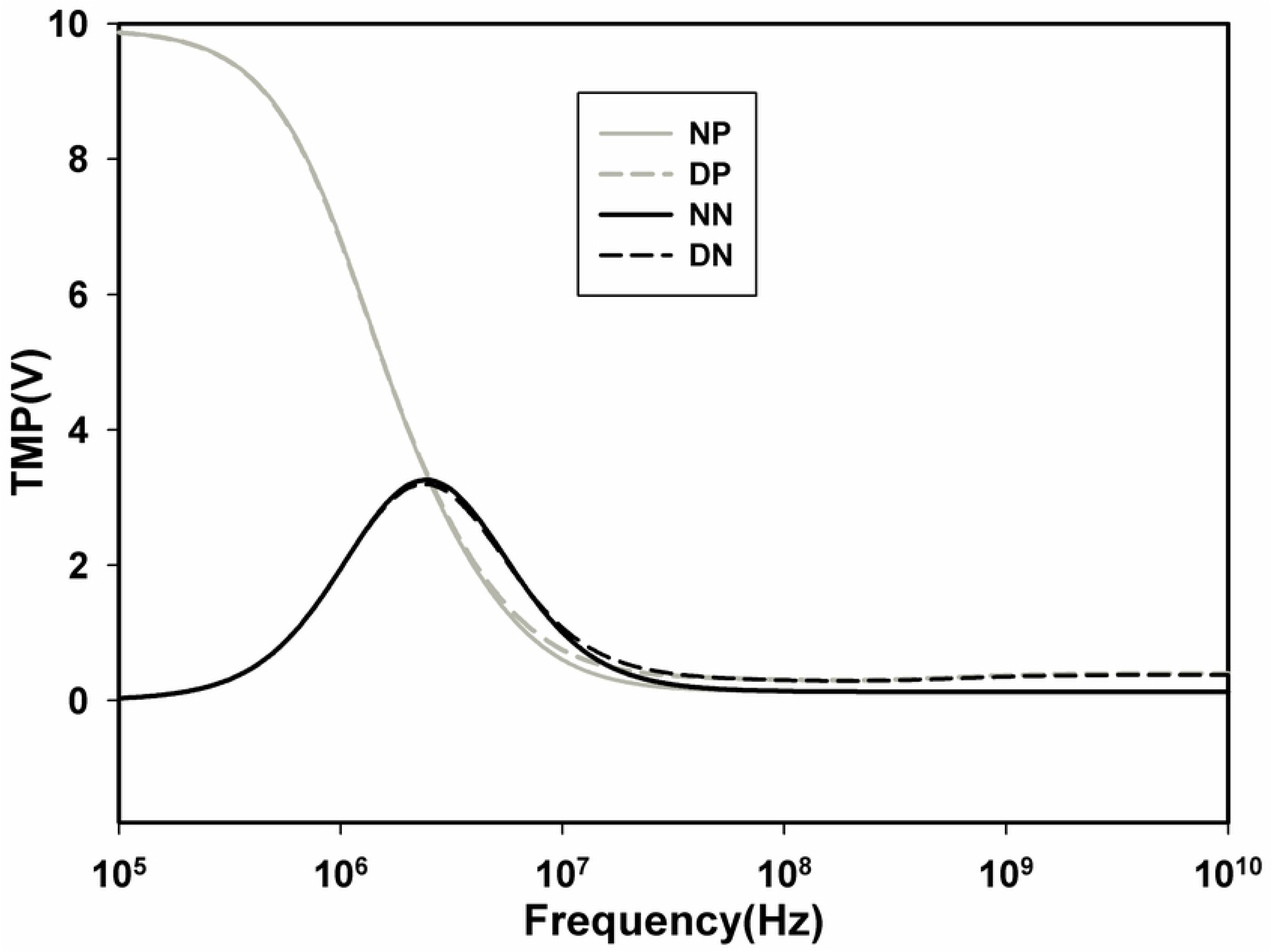
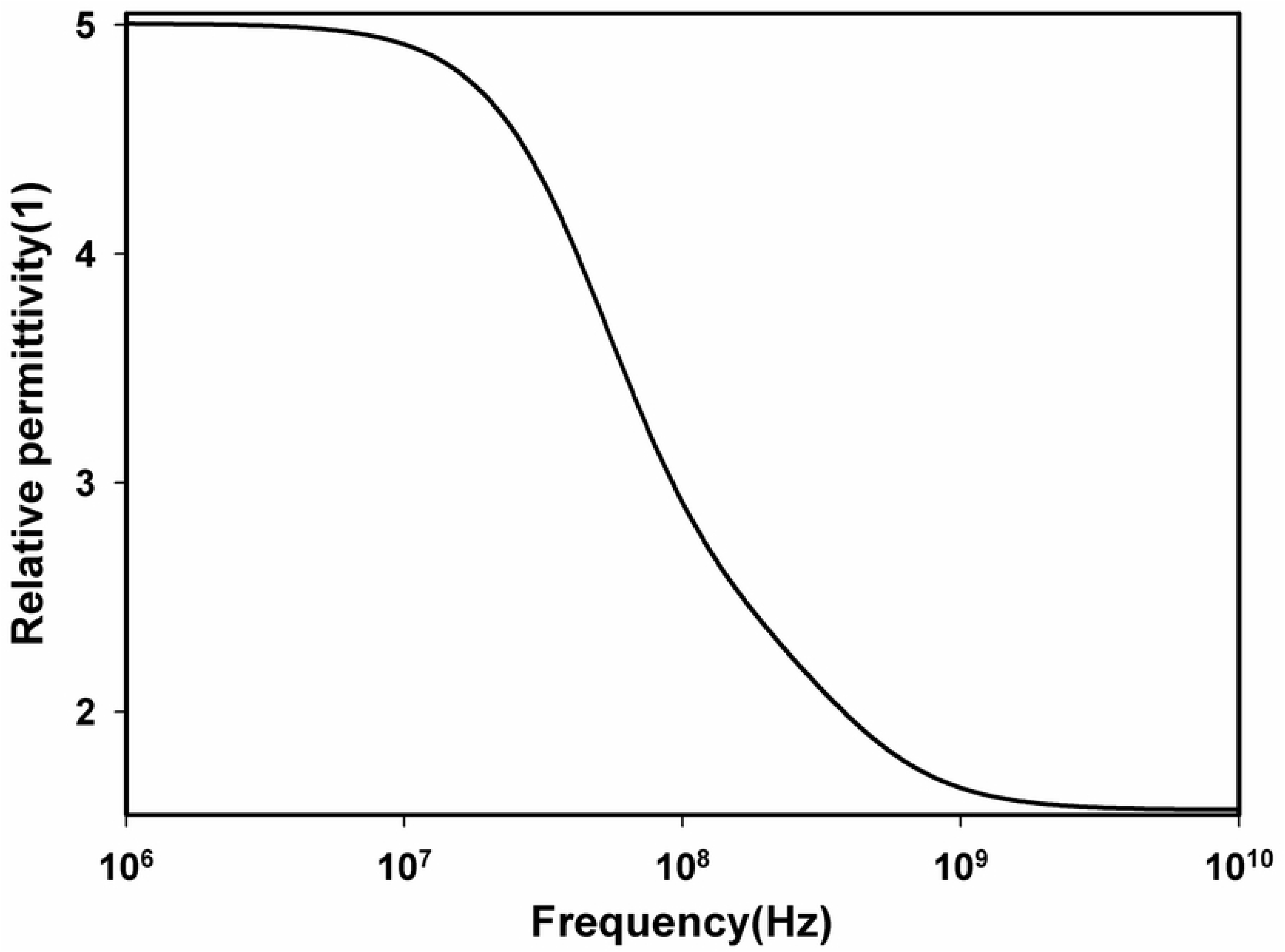
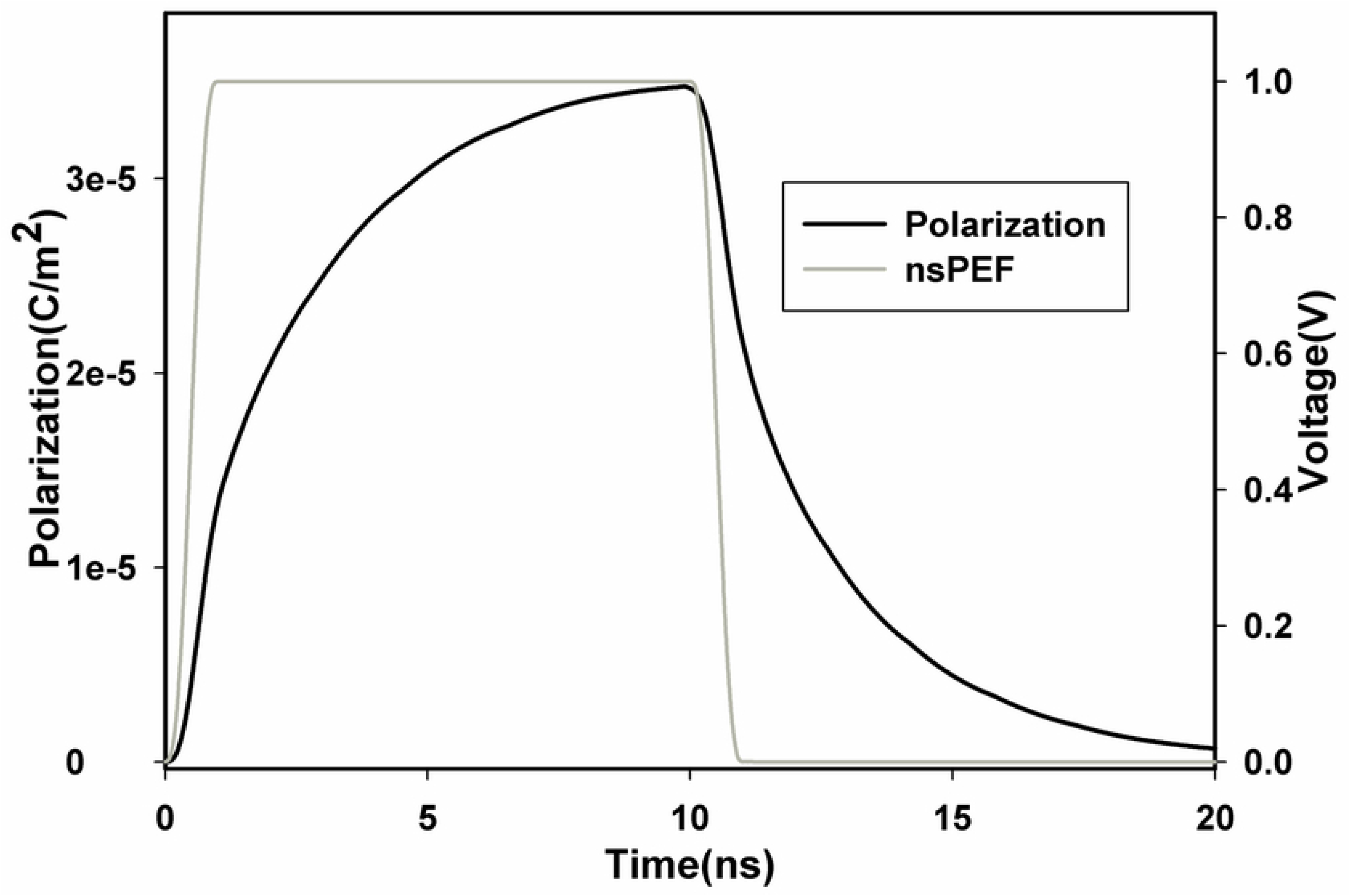
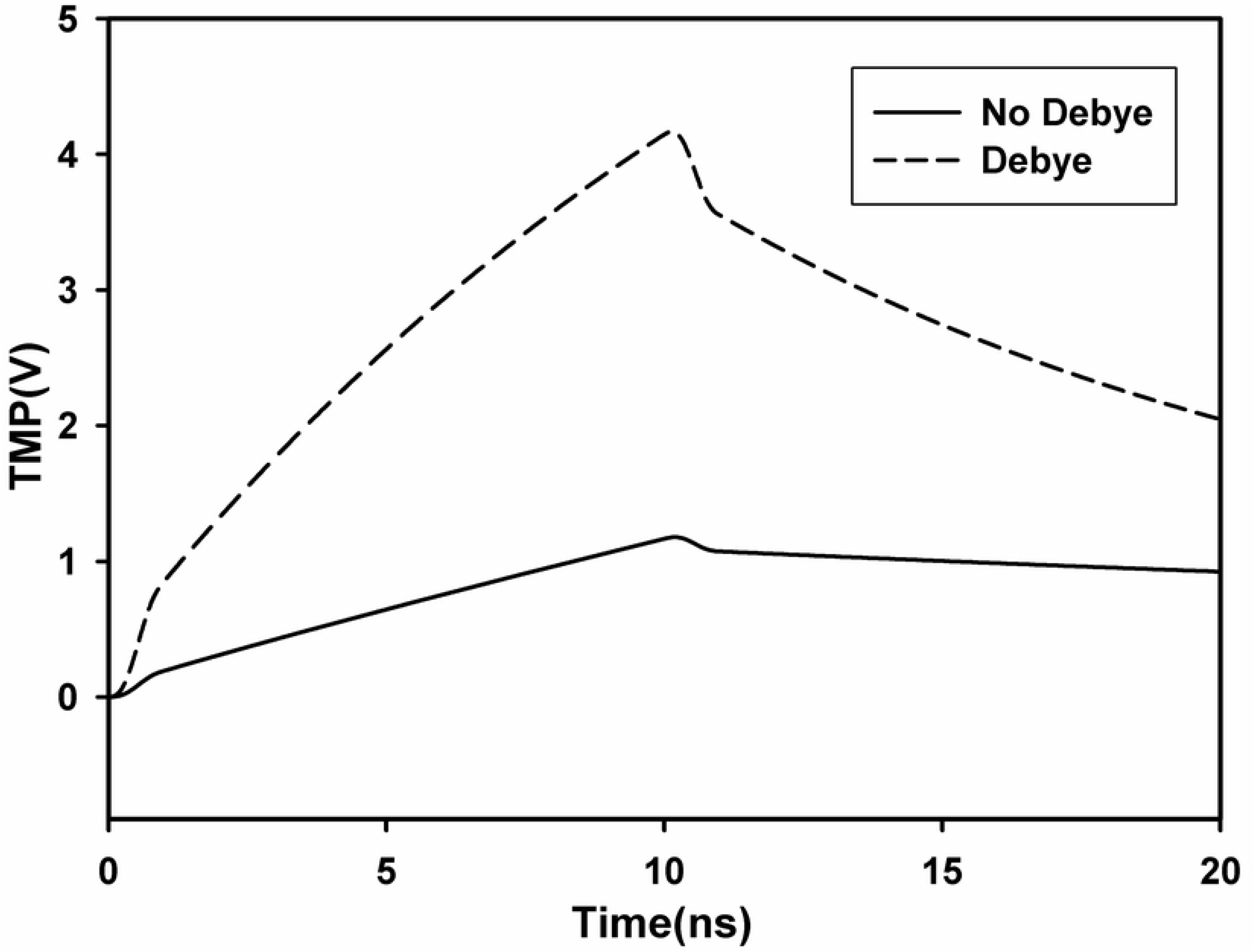
TMP distribution with and without dielectric relaxation. The induced TMP on the cellular membrane (NP for non-dispersive plasma membrane, DP for dispersive plasma membrane) and nuclear membrane (NN for non-dispersive nuclear membrane, DN for dispersive nuclear membrane) versus frequency when the amplitude of the electric field is 10 kV/cm (a), relative permittivity of plasma membrane versus frequency (b), time courses of nsPEFs (gray) and polarization of point A_1_ (black) (c), time courses of TMP of points A1 in dispersive (dotted line) and non-dispersive (solid line) mode (d).

TMP distribution calculated with DP was compared with those without DP, and it indicated that TMP was underestimated from 10^6.5^ to 10^10^ Hz when DP was not taken into account. The relative permittivity of plasma membrane changes with frequency when DP is taken into account in Fig 4b, and the trend is similar to TMP of the plasma membrane.

With the definition of polarization vector P, two-order Debye equation which describes dielectric relaxation of plasma membrane and nuclear membrane in the frequency domain was transformed into the time domain by Laplace transform, then TMP distribution of cell model which includes dielectric relaxation with the application of nsPEFs can be solved in the time domain. The time course of polarization vector of point A_1_ with the application of nsPEFs (pulse duration of 10ns, filed intensity of 10 kV/cm, rise time of 1 ns) is illustrated in Fig 4c. The time trend of polarization vector is almost the same as the nsPEFs except a slower change both during the rising and decreasing periods. The flat top of the polarization vector is about 3.5×10^-5^ C/m^2^, which corresponds to a relative permittivity of 5 (equals to static relative permittivity of plasma membrane), which can prove the correctness of our simulation.

The time course of TMP of point A_1_ with and without DP is shown in Fig 4d, TMP of plasma membrane is always larger with DP than those without during limited observation time, and the biggest difference is about 3 V, furthermore, significant decrease in both the rising and falling periods can be found with DP. The simulation results are in well agreement with previous studies [22–24], which indicate that TMP is underestimated when the DP was not taken into account, in other words, temporal and spatial distribution of TMP can be obtained more accurately with the consideration of dielectric relaxation of all cell compartments.

### C. Temporal and spatial results with both EP and DP

In order to investigate the effects of both DP and EP on the temporal and spatial distribution of TMP of plasma membrane, four nsPEFs with various pulse duration, field intensity and polarity were selected, which are of the same power density to obtain comparable results. Time evolution of TMP and pore density of A_1_ with the application of the above four different nsPEFs in two different modes (EP and DP+EP) is shown in Fig 5, TMP of A_1_ exceeded the critical threshold (1V) with the application of 10ns and 5ns unipolar pulses, however, only the latter pulse induced profound increase in pore density of A_1_, which reached the electroporation threshold (PT=10^15^), in the EP mode.

**Fig 5.**
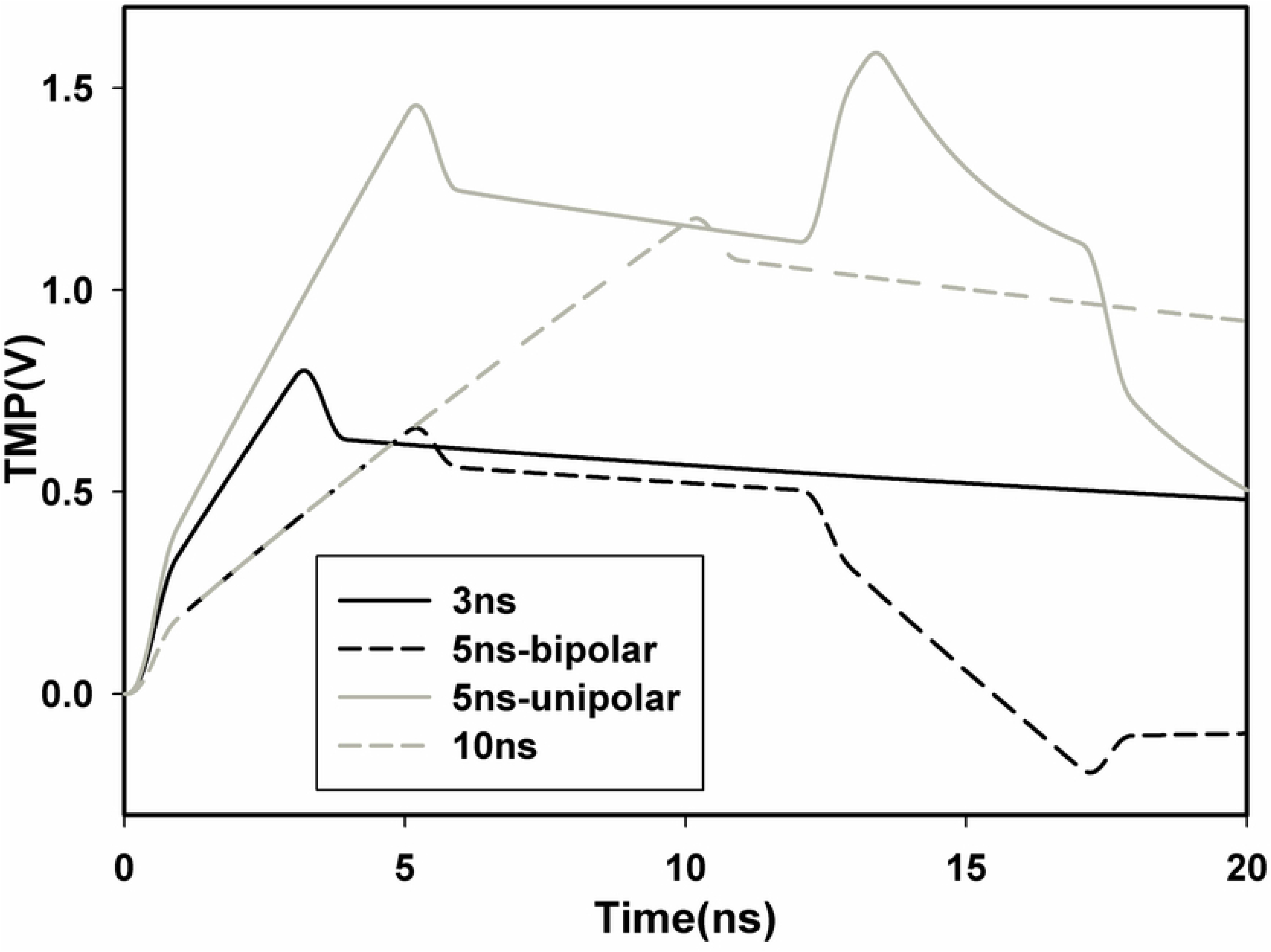
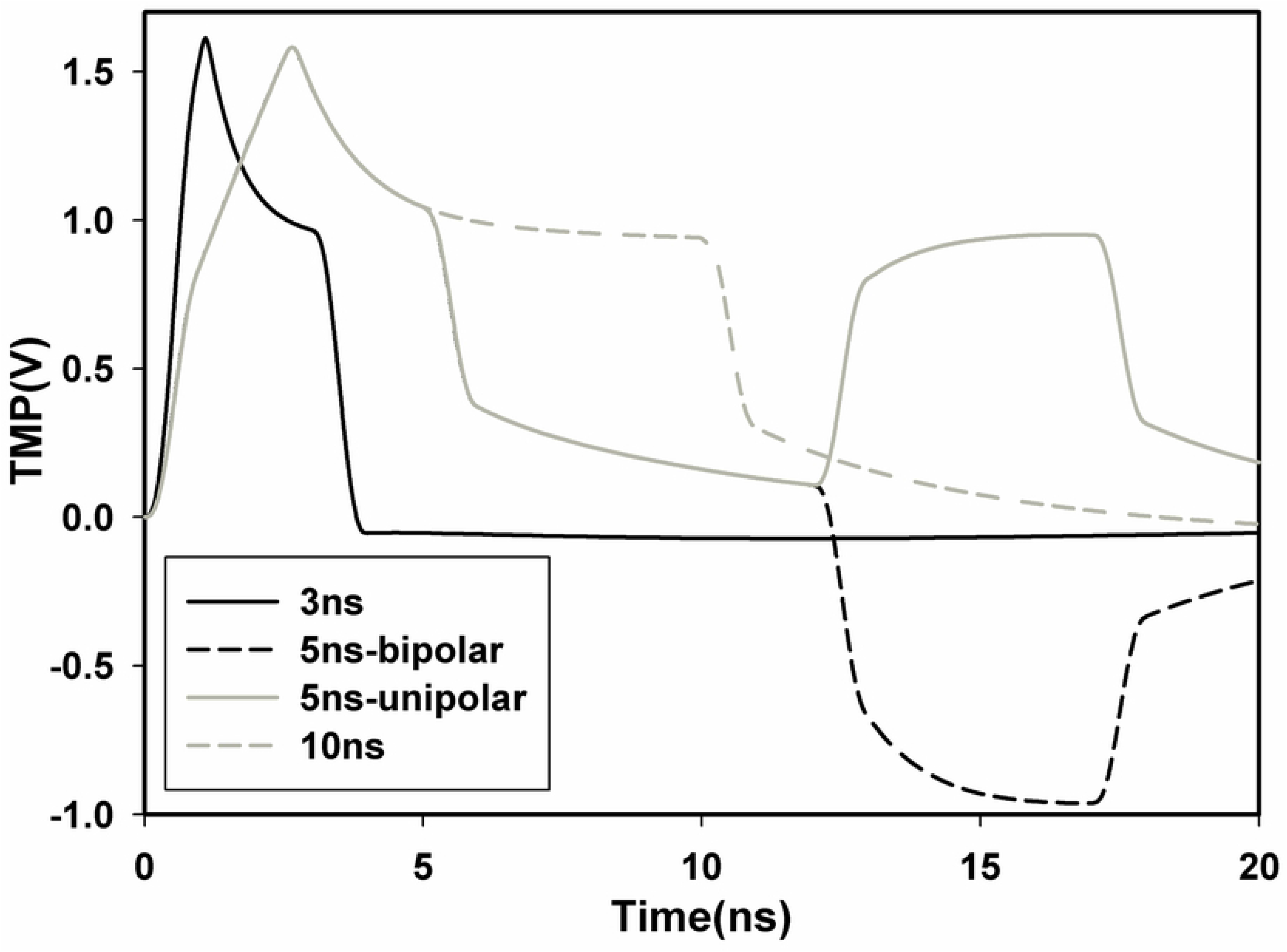
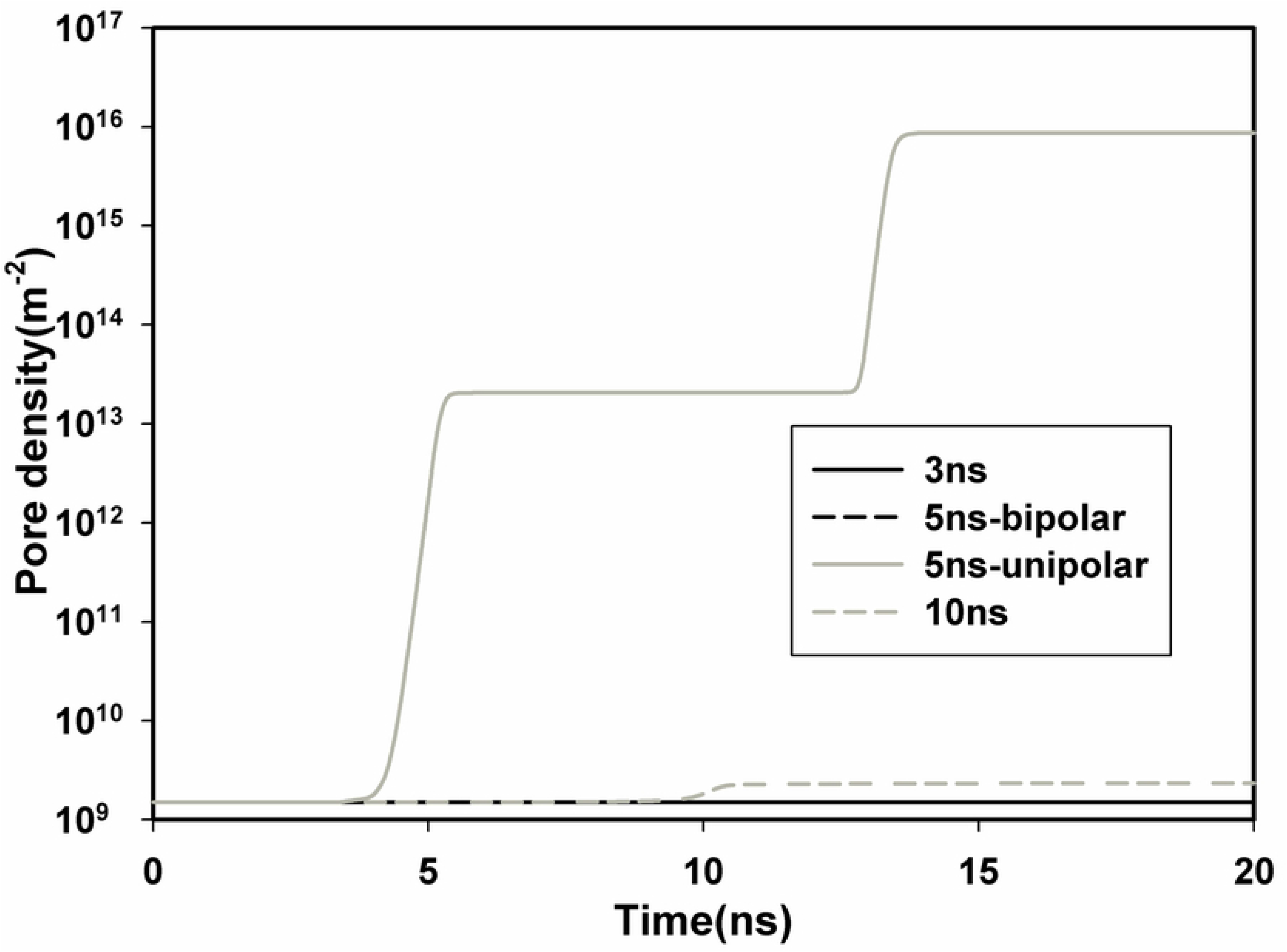
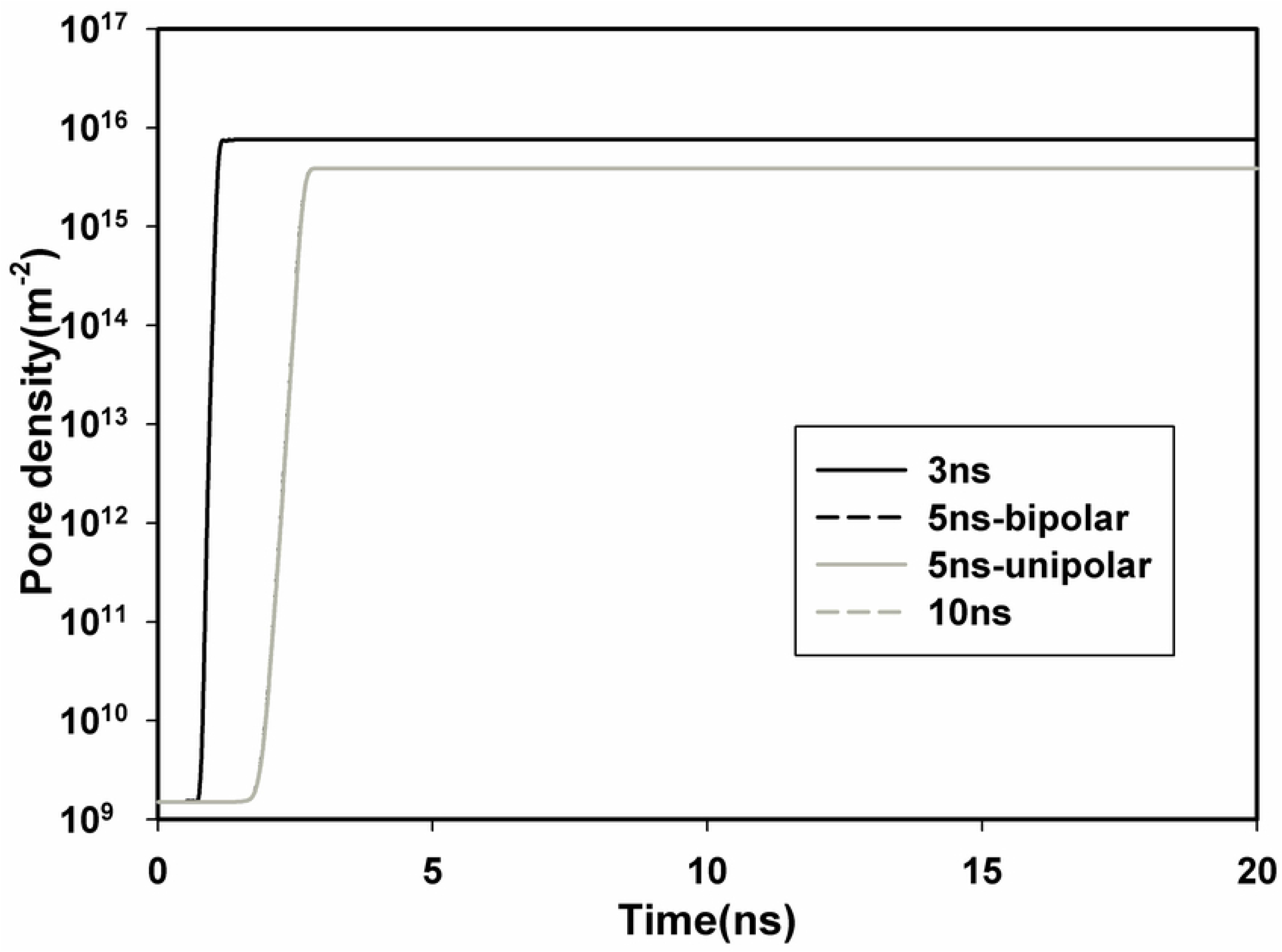
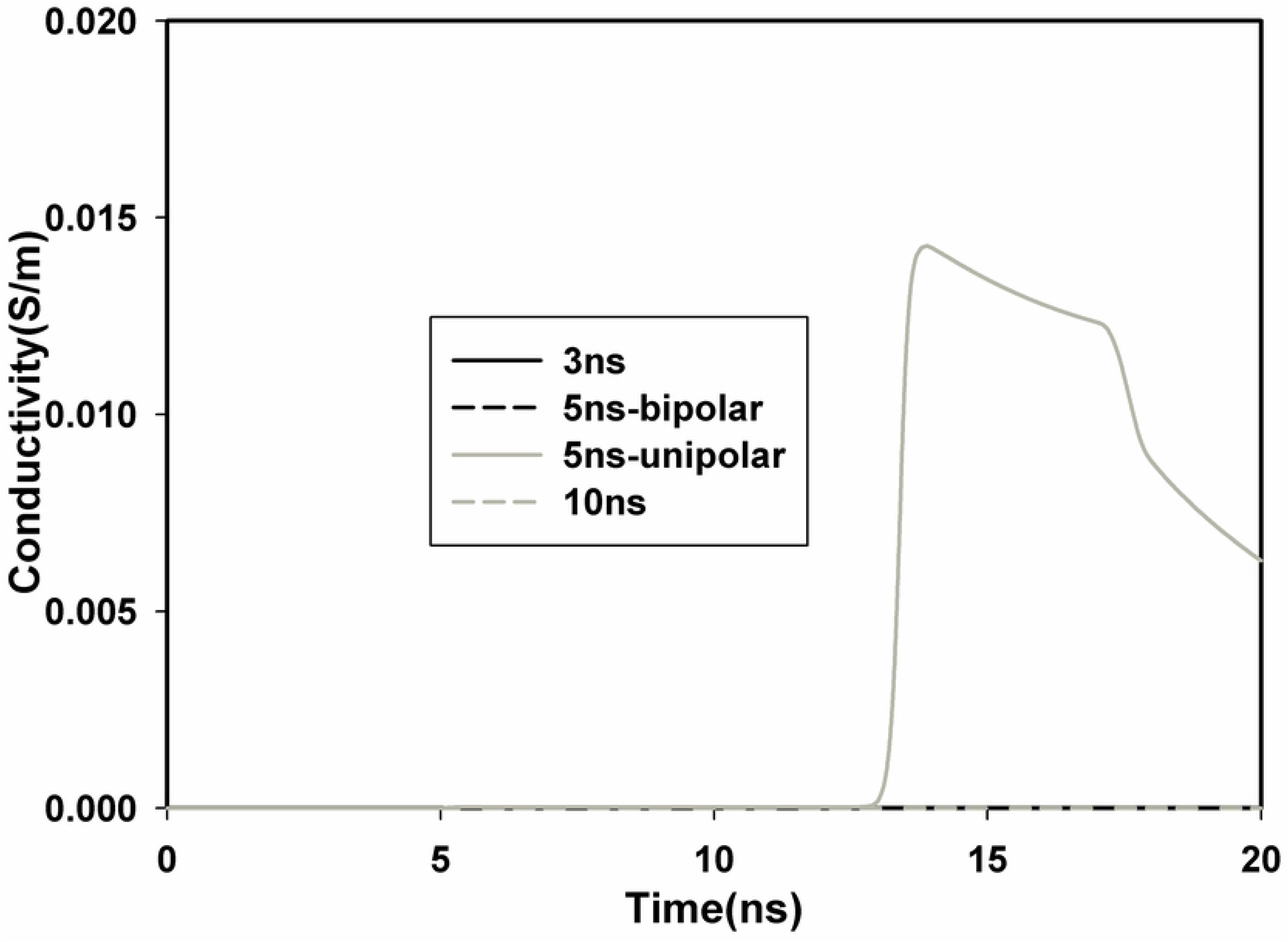
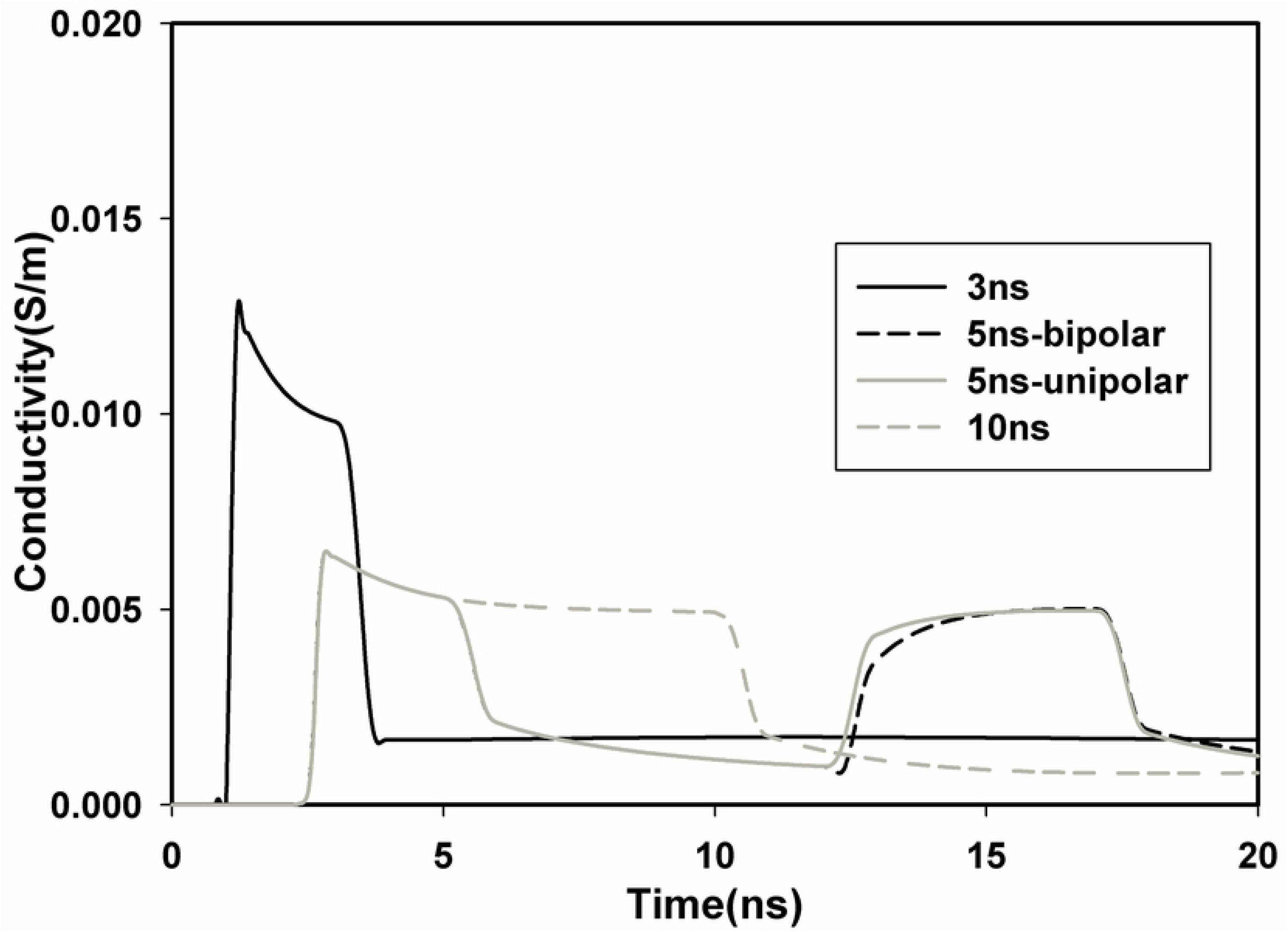
Time evolution of TMP and pore density of A_1_ with four different nsPEFs. Temporal distribution of TMP of A_1_ in EP mode (a) and EP+DP mode (b), pore density of A_1_ in EP mode (c) and EP+DP mode (d), conductivity of A_1_ in EP mode (e) and EP+DP mode (f), with four nsPEFs.

TMP and pore density of A_1_ attained its threshold with all four nsPEFs in the DP+EP mode, and the time required to attain the threshold is much shorter than that of the EP mode, in agreement with [22–24].

After the electroporation threshold PT was overcome the conductivity started to increase, and significant increase in conductivity of A_1_ was observed with only the 5 ns unipolar pulse in EP mode, while significant increase in conductivity of A_1_ was observed with all four nsPEFs in the DP+EP mode.

To get in-depth understanding of the effects of both EP and DP on the temporal and spatial distribution of TMP, we selected seven points separated by 15° in the upper left quarter of the plasma membrane to study the time course of TMP and pore density with the 10 ns pulse, and spatial distribution of TMP and pore density was achieved along the half arc length of plasma membrane from A_1_ to A_8_, both in two different modes (EP and DP+EP). In the EP mode (Figs 6a, c, e and g), the TMP of A_1_ began to increase at 0ns once the pulse was delivered to the cell, reaching a TMP threshold of about 1V at 8.4 ns, then to its peak value of about 1.2 V at 10.2 ns, in agreement with [22]. The time trend is similar in A_2_-A_7_ except a smaller TMP value, and peak values of TMP of A_1_-A_3_ exceed 1V, while A4-A7 is smaller than 1 V. Once the threshold of 1V was overcome the pore density started to increase, in accordance with [22], however, pore density of A_1_ did not reach up to the threshold (PT) of 10^15^ m^-2^ in our simulation, which may due to the differences in model parameters used in our simulation to that of [22]. Spatial distribution of TMP and pore density along the half arc length of plasma membrane gave the similar results, and typical values were listed in Table 1. In addition, significant increase in conductivity of A_1_-A_7_ and along the arc length of plasma membrane was not observed in the EP mode (Figs 6i and k), and the results are in good agreement with the pore density distribution, demonstrating that cell is not effectively electroporated in the EP mode.

**Fig 6.**
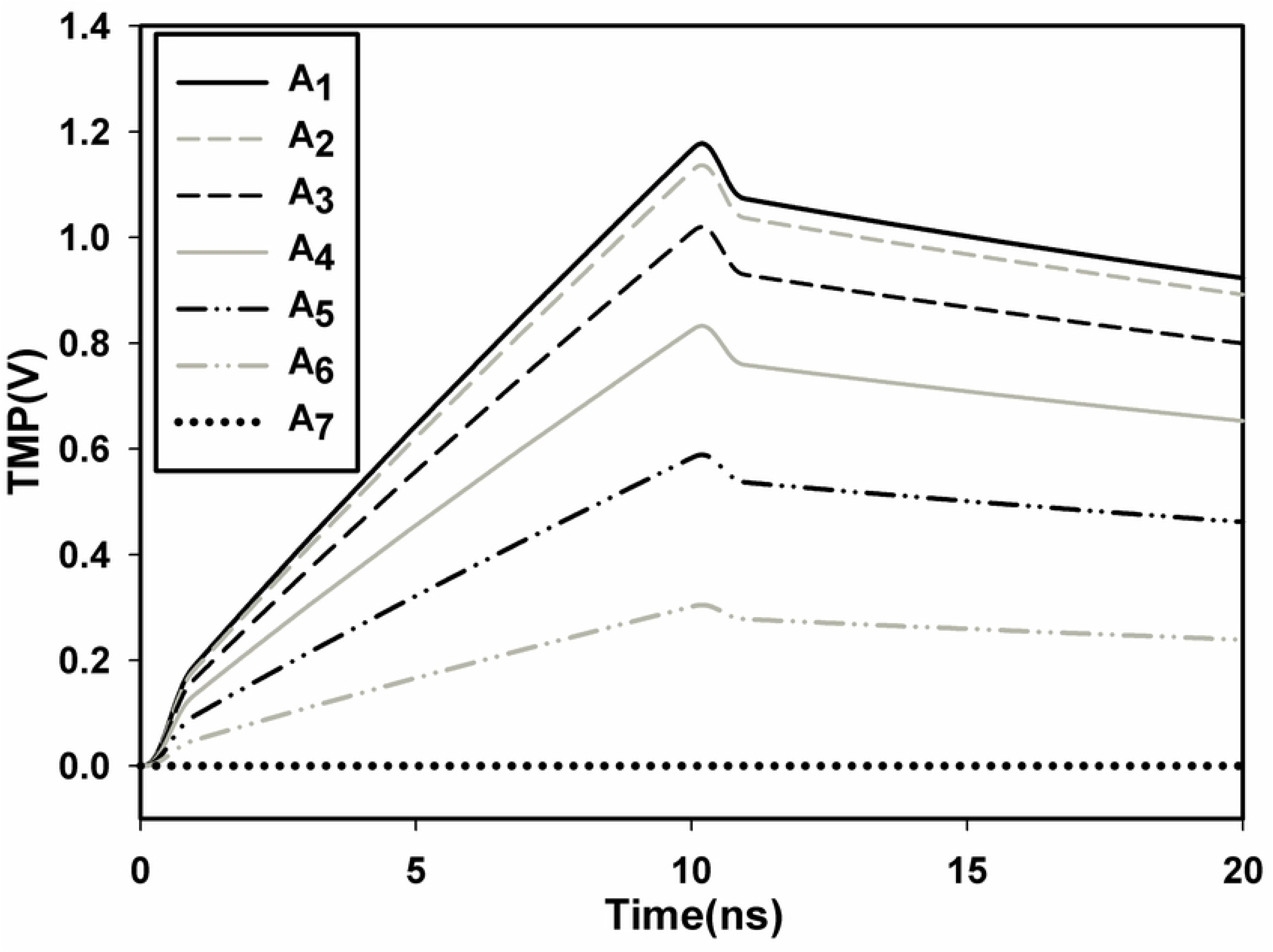
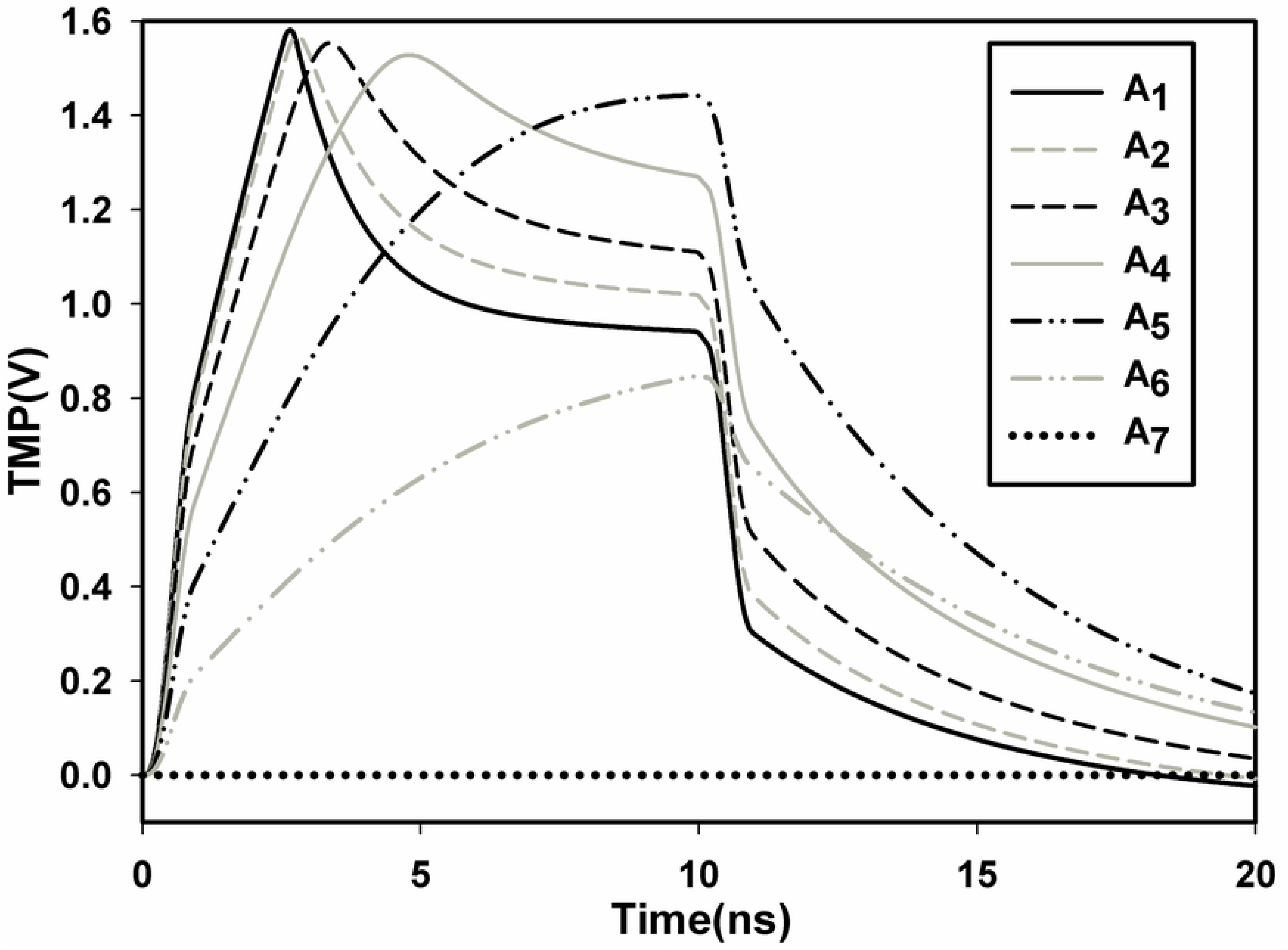
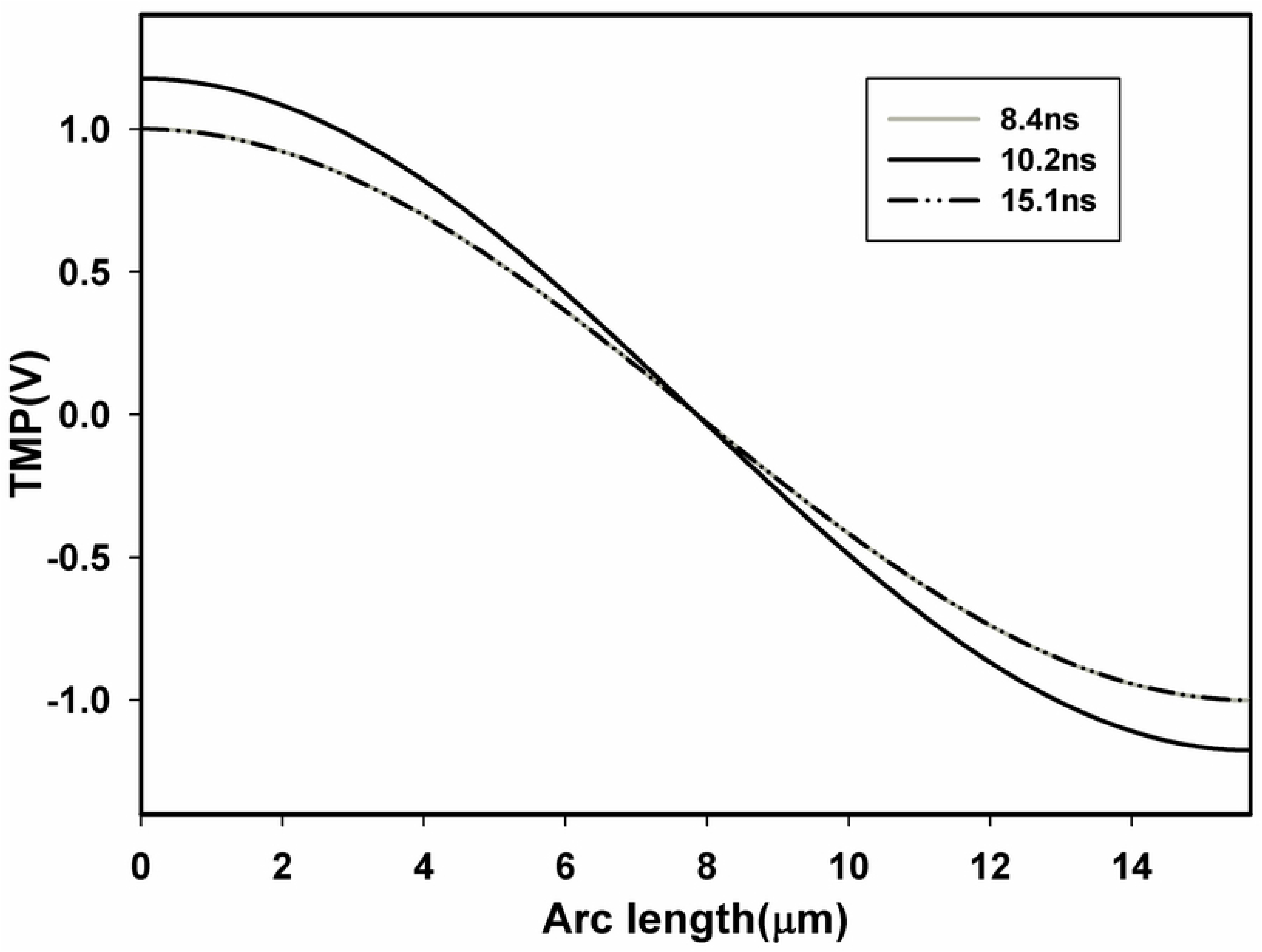
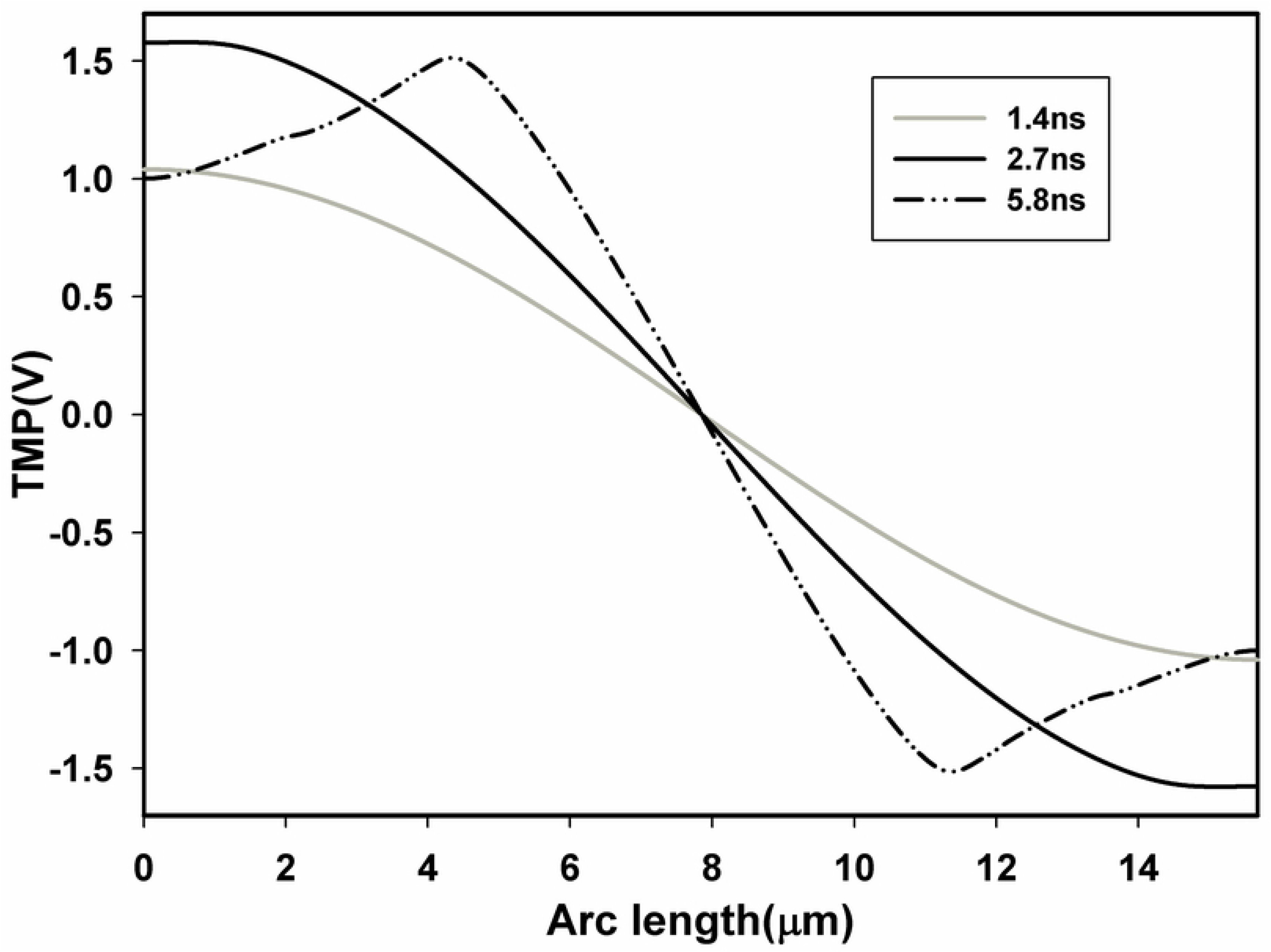
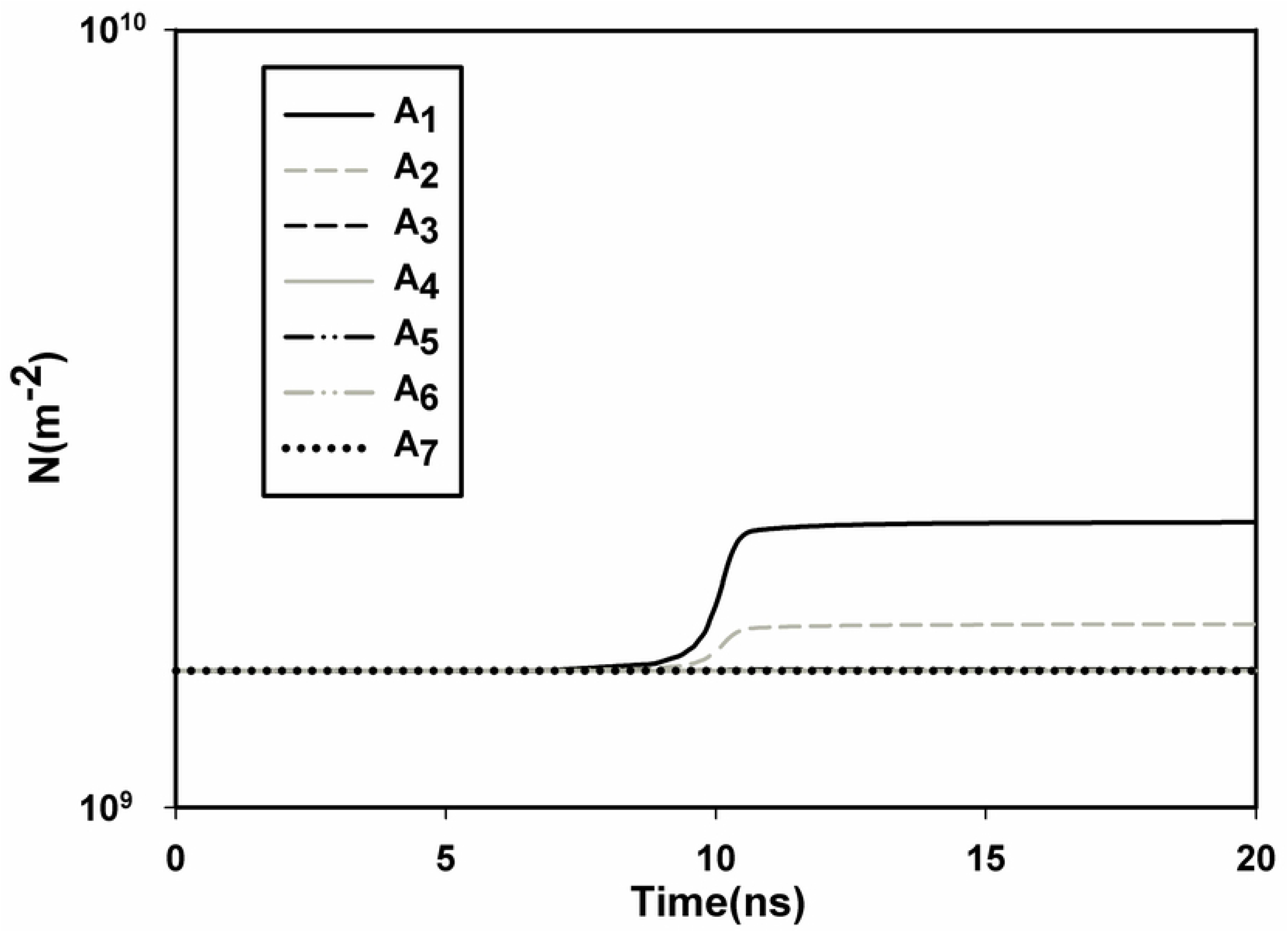
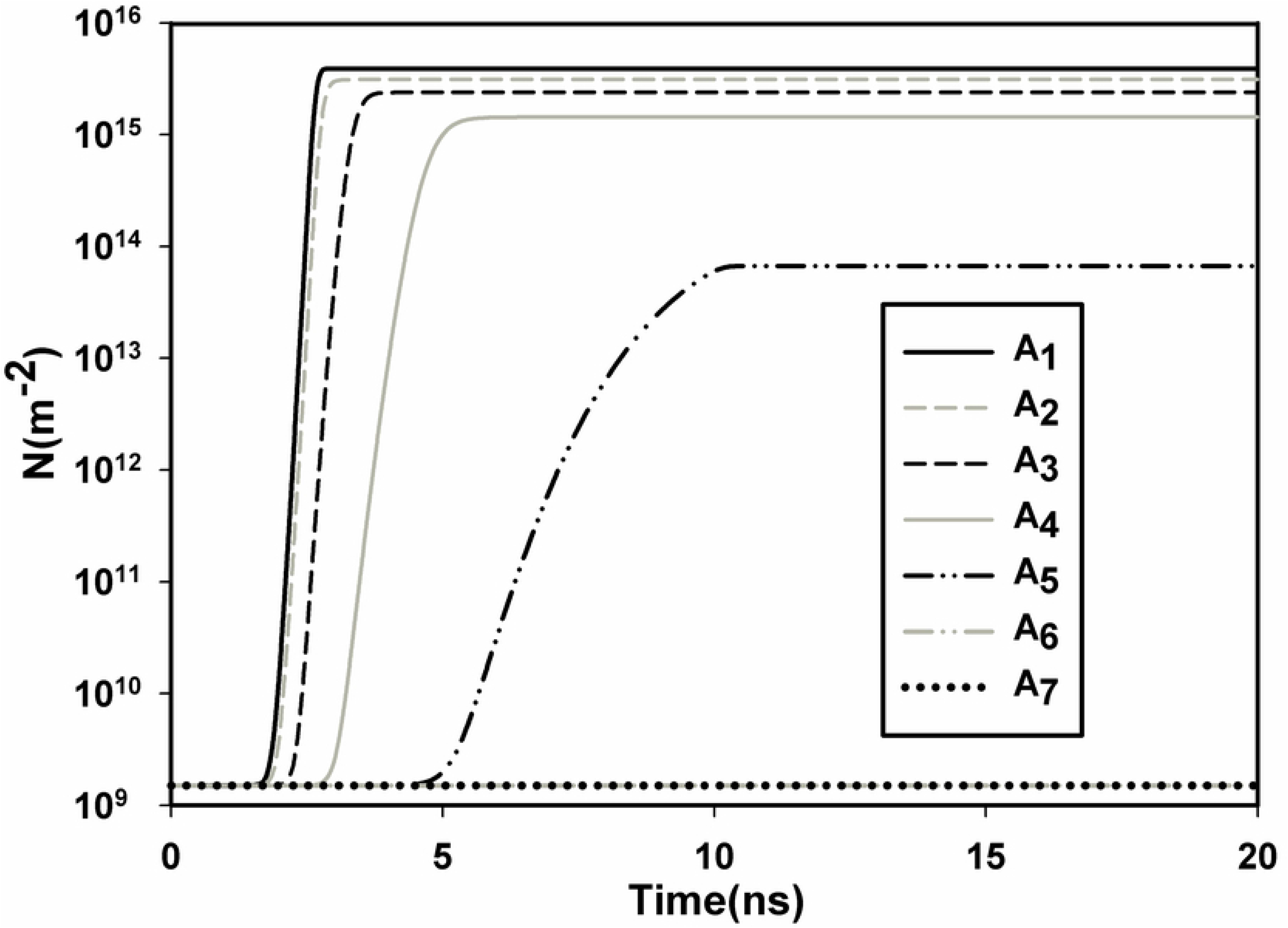
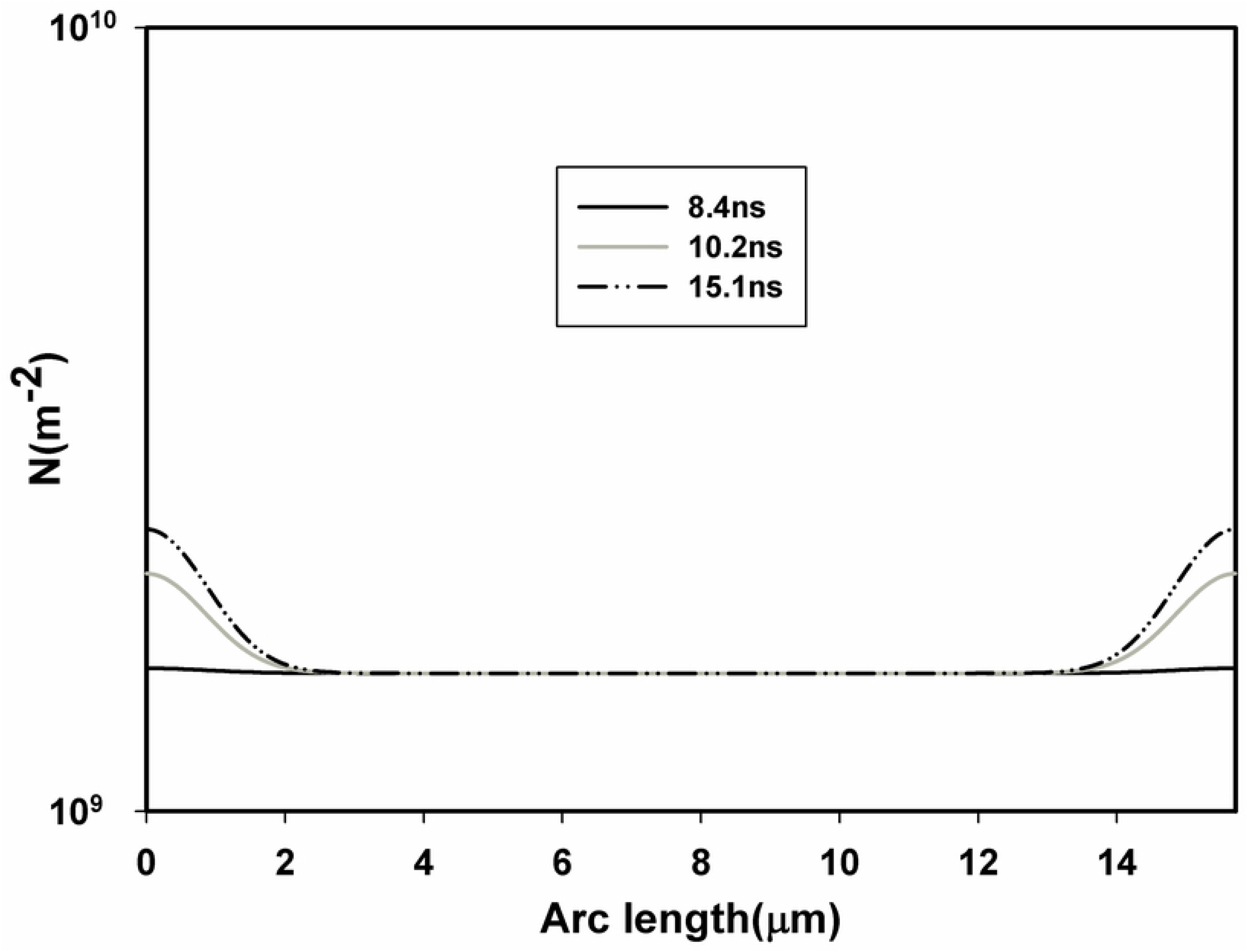
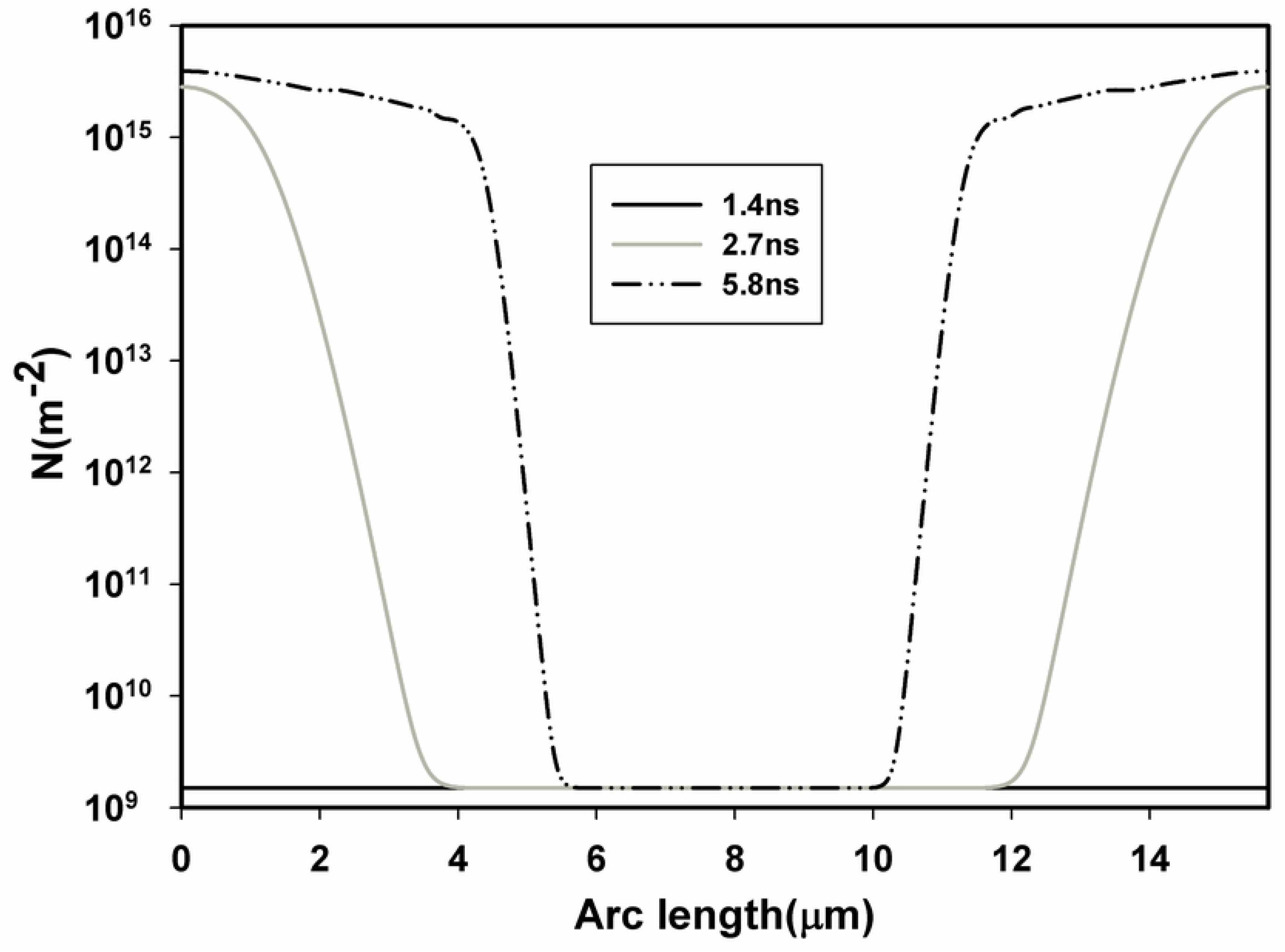
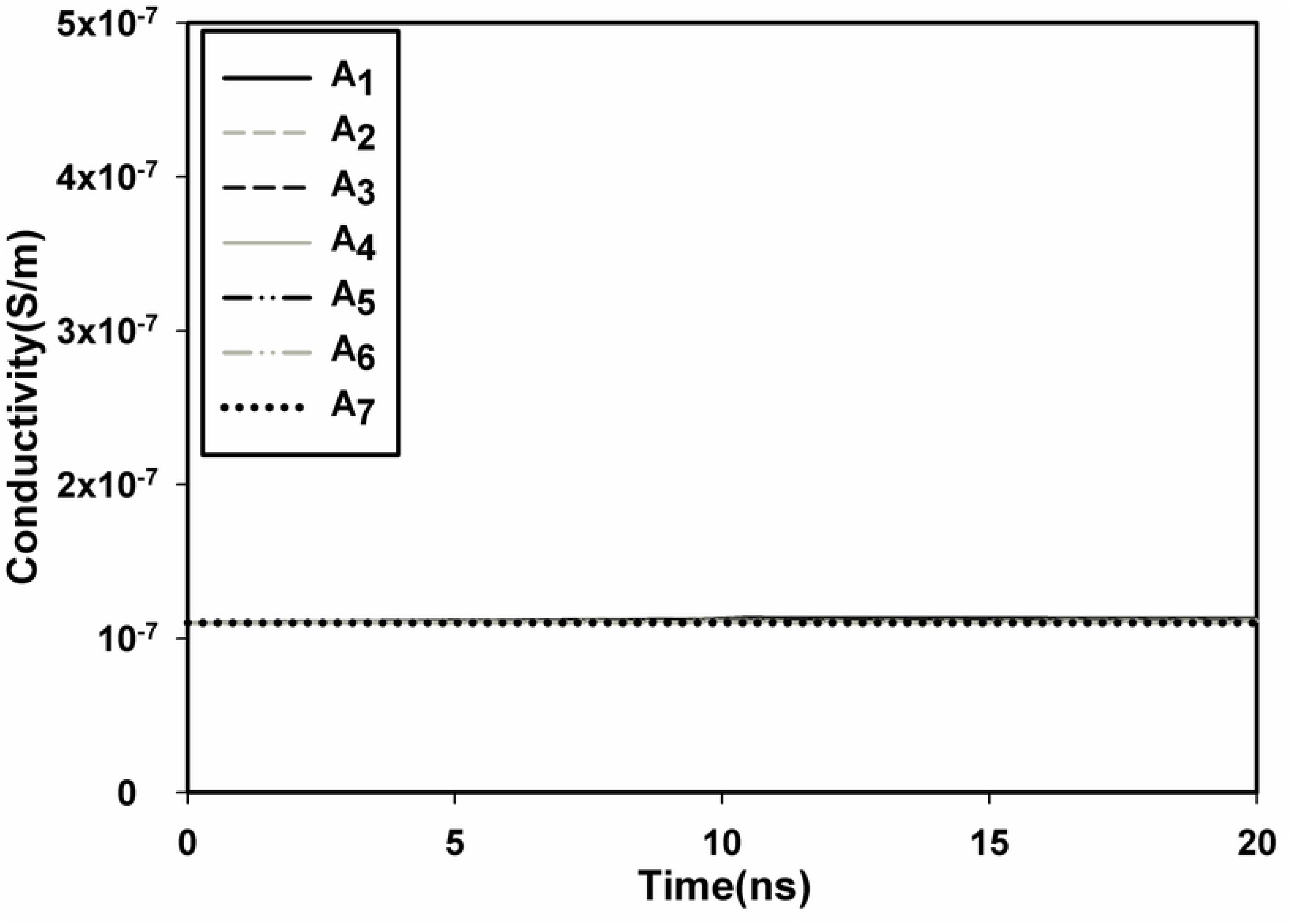
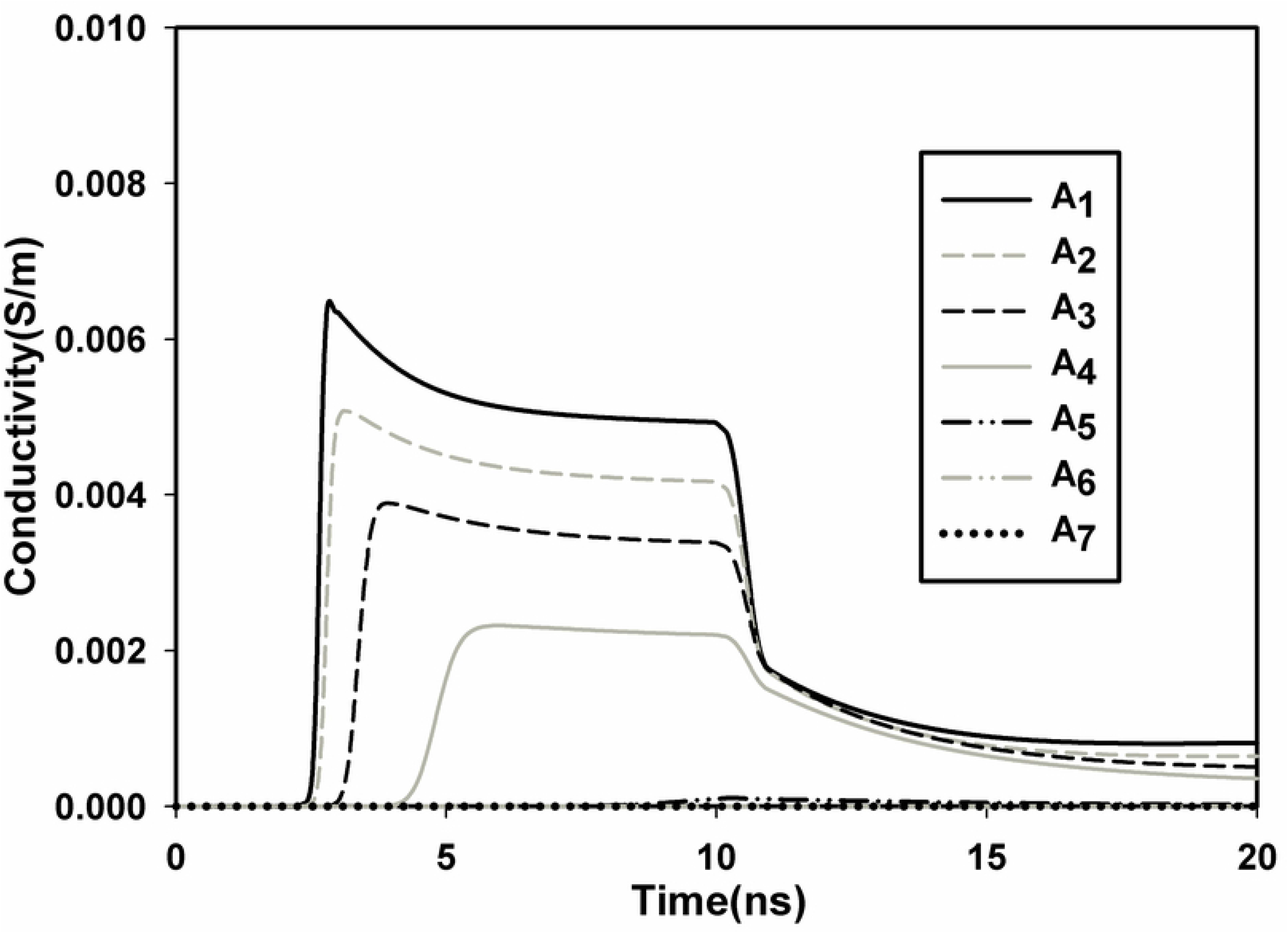
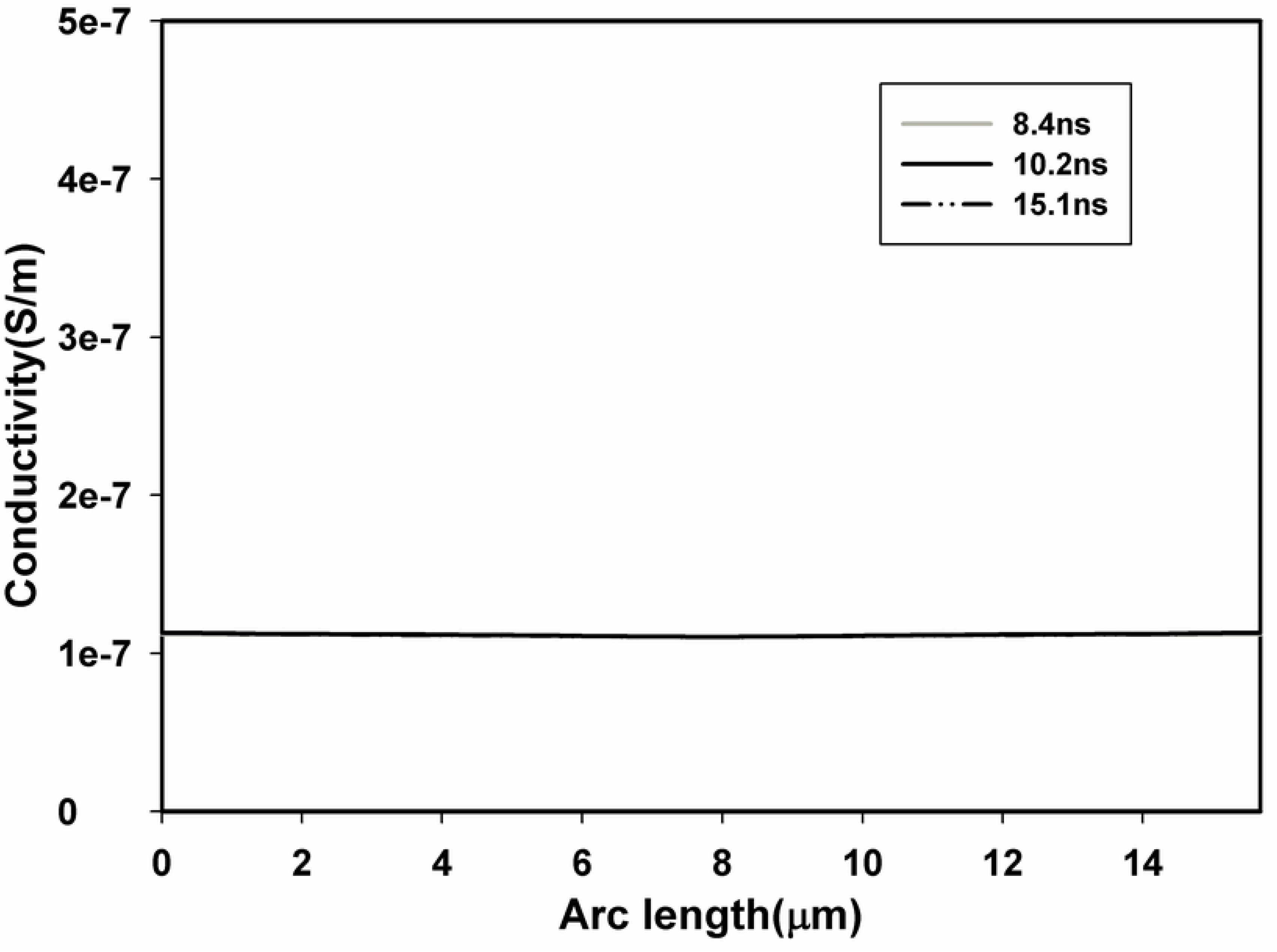
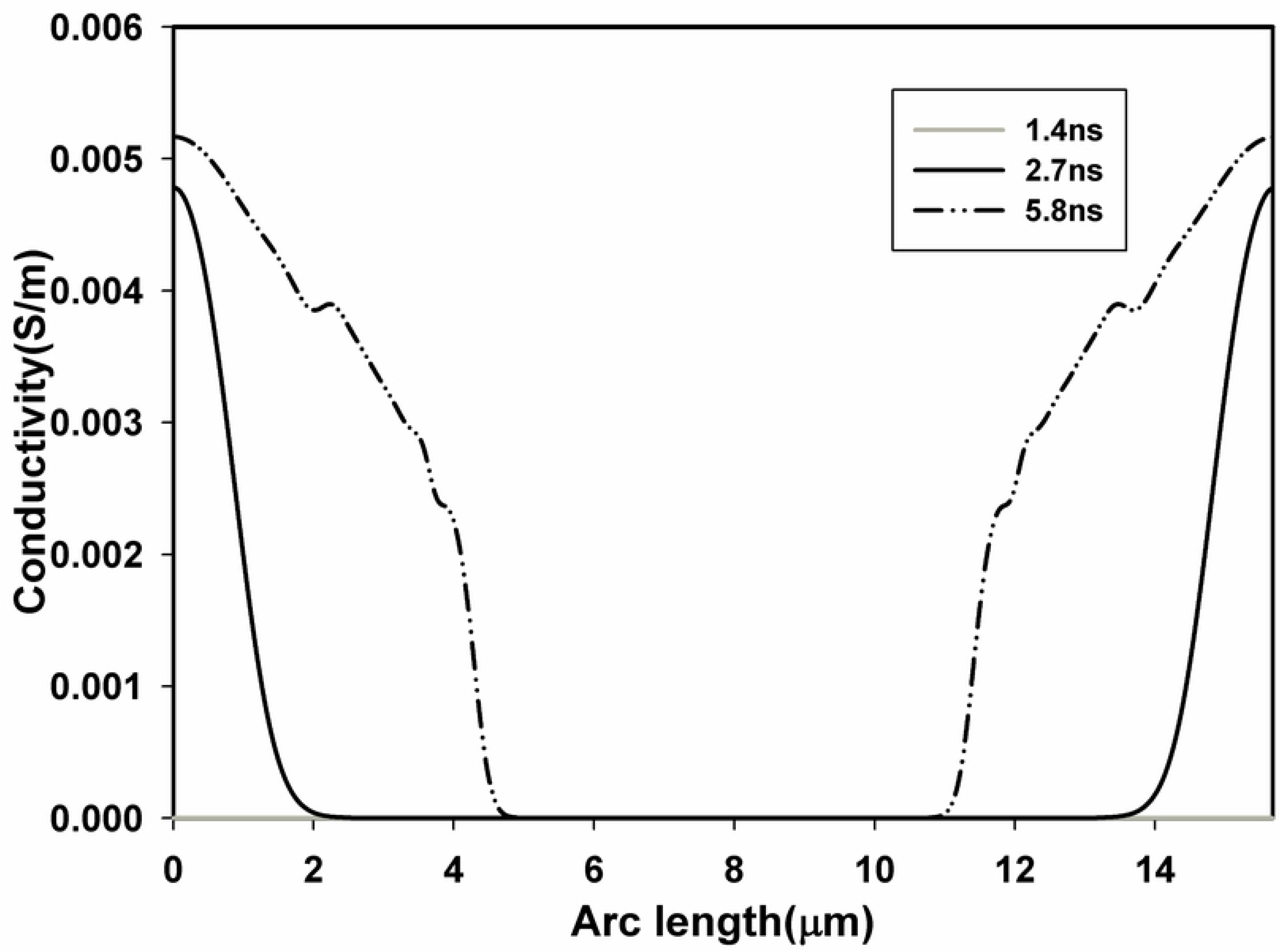
Temporal and spatial results from point A_1_ to A_8_. Temporal distribution of TMP of A_1_-A_7_ in EP mode (a) and EP+DP mode (b), pore density of A_1_-A_7_ in EP mode (e) and EP+DP mode (f), and conductivity of A_1_-A_7_ in EP mode (i) and EP+DP mode (j), spatial distribution of TMP along the arc length of plasma membrane from A_1_ to A_8_ at different times in EP mode (c) and EP+DP mode (d), pore density in EP mode (g)and EP+DP mode (h), and conductivity in EP mode (k) and EP+DP mode (l), when nsPEFs of 10 ns and 10kV/cm is applied.

With the consideration of both EP and DP in the cell model, the TMP of A1 started to increase at 0 ns once the pulse was delivered to the cell, reaching rapidly a TMP threshold of about 1 V at 1.4 ns, then to its peak value of about 1.58 V at 2.7 ns, much faster and larger value of TMP was achieved with the consideration of DP (Figs 6b and d). Once the threshold of 1V was overcome, the pore density began to increase reaching the membrane poration at the threshold of 10^15^ m^-2^, after the PT was overcome the conductivity started to increase reaching about 5 orders of the initial value (Figs 6f, h, j and l), in accordance with [22]. Similar results can be found in A_2_-A_4_ with a decreasing peak value of TMP, flat top value of pore density and conductivity, however, TMP of A_5_ exceeded the threshold of 1 V, while pore density does not overcome the PT and conductivity increased a little bit. Spatial distribution of the TMP, pore density and conductivity along the arc length of plasma membrane gave the similar results, demonstrating that at least 45° near A_1_ of the upper left quarter of the plasma membrane is electroporated, in accordance with [32]. Similar results were also obtained with the application of three other nsPEFs, which is not shown in this paper.

## Discussion and conclusion

Our paper proposes that a microdosimetric study on nsPEFs includes dielectric relaxation of cell plasma membrane and nuclear membrane through a two-order Debye model, and the two-order Debye model is transformed into the time domain with the introduction of polarization vector. Then we obtain the time course of TMP by solving the combination of Laplace equation and time-domain Debye equation. Next, we used the asymptotic version of the smoluchowski equation to characterize electroporation of plasma membrane and added it to our model to predict the temporal and spatial distribution of TMP and pore density.

Our results highlight the relevance of dielectric relaxation in nsPEFs microdosimetry, as evidenced by the fact that both the TMP, pore density and the conductivity are strongly influenced by the dispersion. TMP, pore density and conductivity is underestimated if Debye model is disregarded. Therefore, for pulse duration less than or equal to 10ns, the inclusion of the Debye equation in the characterization of the cell compartments is necessary to accurately quantify TMP, pore density and conductivity.

During the evaluation of this simulation, we noted that it was unable to find all of the parameters for a single cell in literature. The parameters listed in Table.1, such as cell geometrical size, conductivity and permittivity of all components, were obtained from external sources, other theoretical models, or experiments. Thus, differences between experimental results and simulation results are predictable. In order to prove the correctness of our simulation, we evaluated the time course of TMP of A_1_, and compared the simulation results with that of the analytical results obtained with the first-order Schwan equation, both used the same model parameters listed in Table.1, and our algorithm gives satisfactory accuracy with a maximum difference of about 2%. TMP distribution both in the frequency and time domain is underestimated without considering dielectric relaxation during specific frequency or with pulse duration less than or equal to 10 ns, and this trend is in well agreement with previous studies, furthermore, correctness of the interpretation of Debye model in frequency and the time domain can be proved by the spectrum distribution of relative permittivity and time course of the polarization vector.

One unique aspect of this study is to include both DP and EP in the dielectric double-shelled cell model, to obtain the temporal and spatial distribution of TMP of plasma membrane without the introduction of complex mathematics. And the algorithm presented in this study can be easily applied to biological cell of irregular shapes, even to real biological cells.

In EP mode, TMP of A_1_-A_7_ follows the cosθ law, as evidenced by the peak value of TMP listed in Table.1, which means that plasma membrane is not electroporated, as previous experimental studies demonstrated that the cosθ law is not valid once significant poration occurs, and the results are in accordance with the pore density and conductivity distribution, where pore density electroporation threshold PT is not overcome and no significant increase in conductivity is observed. In EP+DP mode (Table.2), TMP of different points on plasma membrane does not follow the cosθ law, and pore density electroporation threshold PT is overcome in A_1_-A_4_, where significant increase in conductivity is also found, demonstrate that at least 45° of 90° of plasma membrane is electroporated. Krassowska and Filev [32] found that the boundary of the electroporation and the nonelectroporation is 45°, and this value is similar to our simulation results.

In addition, Fig 6f shows that the location on the membrane closest to the electrodes has the largest pore density, and the pore density decreases from the point to the pole. Pucihar and colleagues [27] observed that the electrode near the membrane had the maximum fluorescence intensity, which was consistent with our results. Significant increase of about 5 orders was observed in the conductivity of A_1_-A_4_ in Fig 6j, in agreement with [33], in which conductivity of an oxidized cholesterol membrane with the application of 20 μs pulse was measured, and significant increase of 4 to 5 orders in conductivity was found. Although previous studies showed that nsPEFs induced more pronounced increase in conductivity through EP than that of μsPEFs [18], our simulation can give comparable results.

**Table 2.** Typical values obtained from Fig 6, which includes peak value of TMP and flat top value of pore density at different points, in both EP and DP+EP modes, and the time required to attain the typical values is also taken into account. Commonly, TMP of 1 V and pore density of 10^15^ is used as threshold value to predict the appearance of electroporation.

| Points | Peak value of TMP (EP)/time required to reach the peak value | Peak value of TMP(EP+DP)/time required to reach the peak value | Flat top value of pore density (EP) | Flat top value of pore density (EP+DP) |
| --- | --- | --- | --- | --- |
| A<sub>1</sub> | 1.18 V/10.2 ns | 1.58 V/2.7 ns | 2.3e9/10.7 ns | 3.90e15/3.1 ns |
| A<sub>2</sub> | 1.14 V/10.2 ns | 1.57 V/2.8 ns | 1.7e9/10.7 ns | 3.11e15/3.4 ns |
| A<sub>3</sub> | 1.02 V/10.2 ns | 1.55 V/3.4 ns | N<sub>0</sub> | 2.39e15/4.1 ns |
| A<sub>4</sub> | 0.83 V/10.2 ns | 1.53 V/4.8 ns | N<sub>0</sub> | 1.44e15/6.6 ns |
| A<sub>5</sub> | 0.59 V/10.2 ns | 1.44 V/9.9 ns | N<sub>0</sub> | 6.67e13/10.5 ns |
| A<sub>6</sub> | 0.30 V/10.2 ns | 0.85 V/10 ns | N<sub>0</sub> | N<sub>0</sub> |
| A<sub>7</sub> | 0 V | 0 | N<sub>0</sub> | N<sub>0</sub> |

In EP mode, TMP of 3 ns and 5 ns bipolar pulses did not reach TMP threshold of 1 V, while TMP of 10 ns and 5 ns unipolar pulses reached 1 V, however, cell was electroporated with only the application 5 ns unipolar pulse, as evidenced by the fact that significant increase in pore density and conductivity was observed with only the 5 ns unipolar pulse, which means that only TMP threshold of 1 V is not sufficient to predict the EP of biological cell, time evolution of pore density and (or) conductivity need to be taken into account.

To conclude, our results demonstrate that performing nsPEFs dosimetry at the single cell level is useful to accurately predict the temporal and spatial distribution of TMP, pore density and conductivity. This type of predictive analysis is effective for optimizing in the use of pulse generators and applicators in terms of the pulse amplitude and waveform for medical application needed of the nsPEFs.

In this study, only dielectric relaxation of plasma membrane and nuclear membrane were included, however, dielectric relaxation of the extracellular medium and cytoplasm has to be included when spectrum of PEF exceeds 20 GHz [21]. The pore radius which was considered constant in our study varies with time and space and need to be considered in more detailed model [32]. Furthermore, biological cells with irregular shape or real cells should be modelled instead of a spherical cell.

## Acknowledgment

This work was supported in part by the National Natural Science Foundation of China (51507024), in part by the innovation Team Program of Chongqing Education Committee (CXTDX201601019).

## References

1. Neumann ES, A. E. Sowers, C. A. Jordan, Eds. (1989), Electroporation and Electrofusion in Cell Biology. New York, NY, USA: Plenum.

2. Ho, S. Y., & Mittal, G. S. (1996). Electroporation of cell membranes: a review. Critical reviews in biotechnology, 16(4), 349–362. https://doi.org/10.3109/07388559609147426

3. Okino, M., & Mohri, H. (1987). Effects of a high-voltage electrical impulse and an anticancer drug on in vivo growing tumors. Japanese Journal of Cancer Research GANN, 78(12), 1319–1321. https://doi.org/10.20772/cancersci1985.78.12_1319 PMID:2448275

4. Tekle, E., Astumian, R. D., & Chock, P. B. (1991). Electroporation by using bipolar oscillating electric field: an improved method for DNA transfection of NIH 3T3 cells. Proceedings of the National Academy of Sciences, 88(10), 4230–4234. https://doi.org/10.1073/pnas.88.10.4230 PMID: 2034667

5. Mir, L. M., Orlowski, S., Belehradek Jr, J., Teissie, J., Rols, M. P., Serša, G., … & Heller, R. (1995). Biomedical applications of electric pulses with special emphasis on antitumor electrochemotherapy. Bioelectrochemistry and Bioenergetics, 38(1), 203–207. https://doi.org/10.1016/0302-4598(95)01823-W

6. Heller, R., Coppola, D., Pottinger, C., Gilbert, R., & Jaroszeski, M.J. (2002). Effect of electrochemotherapy on muscle and skin. Technology in cancer research & treatment, 1(5), 385–391. https://doi.org/10.1177/153303460200100509 PMID: 12625764

7. Schoenbach, K. H., Peterkin, F. E., Alden, R. W., & Beebe, S.J. (1997). The effect of pulsed electric fields on biological cells: Experiments and applications. IEEE transactions on plasma science, 25(2), 284–292. DOI: 10.1109/27.602501

8. Nuccitelli, R., Pliquett, U., Chen, X., Ford, W., Swanson, R. J., Beebe, S. J., … & Schoenbach, K. H. (2006). Nanosecond pulsed electric fields cause melanomas to self-destruct. Biochemical and biophysical research communications, 343(2), 351–360. https://doi.org/10.1016/j.bbrc.2006.02.181 PMID: 16545779

9. Nuccitelli, R., Chen, X., Pakhomov, A. G., Baldwin, W. H., Sheikh, S., Pomicter, J. L., … & Beebe, S. J. (2009). A new pulsed electric field therapy for melanoma disrupts the tumor’s blood supply and causes complete remission without recurrence. International journal of cancer, 125(2), 438–445. doi: 10.1002/ijc.24345. PMID: 19408306

10. Beebe, S., Sain, N., & Ren, W. (2013). Induction of cell death mechanisms and apoptosis by nanosecond pulsed electric fields (nsPEFs). Cells, 2(1), 136–162. https://doi.org/10.3390/cells2010136 PMID: 24709649

11. Esser, A. T., Smith, K. C., Gowrishankar, T. R., & Weaver, J. C. (2009). Towards solid tumor treatment by nanosecond pulsed electric fields. Technology in cancer research & treatment, 8(4), 289–306. https://doi.org/10.1177/153303460900800406 PMID: 19645522

12. Gowrishankar, T. R., Esser, A. T., Vasilkoski, Z., Smith, K. C., & Weaver, J. C. (2006). Microdosimetry for conventional and supra-electroporation in cells with organelles. Biochemical and biophysical research communications, 341(4), 1266–1276. https://doi.org/10.1016/j.bbrc.2006.01.094 PMID: 16469297

13. Schoenbach, K. H., Beebe, S. J., & Buescher, E. S. (2001). Intracellular effect of ultrashort electrical pulses. Bioelectromagnetics: Journal of the Bioelectromagnetics Society, The Society for Physical Regulation in Biology and Medicine, The European Bioelectromagnetics Association, 22(6), 440–448. PMID: 11536285

14. Tekle, E., Oubrahim, H., Dzekunov, S. M., Kolb, J. F., Schoenbach, K. H., & Chock, P. B. (2005). Selective field effects on intracellular vacuoles and vesicle membranes with nanosecond electric pulses. Biophysical journal, 89(1), 274–284. https://doi.org/10.1529/biophysj.104.054494 PMID: 15821165

15. White, J. A., Blackmore, P. F., Schoenbach, K. H., & Beebe, S. J. (2004). Stimulation of capacitative calcium entry in HL-60 cells by nanosecond pulsed electric fields. Journal of Biological Chemistry, 279(22), 22964–22972. doi: 10.1074/jbc.M311135200 PMID: 15821165

16. Gowrishankar, T. R., & Weaver, J. C. (2006). Electrical behavior and pore accumulation in a multicellular model for conventional and supra-electroporation. Biochemical and biophysical research communications, 349(2), 643–653. https://doi.org/10.1016/j.bbrc.2006.08.097 PMID: 16959217

17. Lamberti, P., Romeo, S., Sannino, A., Zeni, L., & Zeni, O. (2015). The role of pulse repetition rate in nsPEF-induced electroporation: a biological and numerical investigation. IEEE Transactions on Biomedical Engineering, 62(9), 2234–2243. DOI: 10.1109/TBME.2015.2419813 PMID: 25850084

18. Zhuang, J., Ren, W., Jing, Y., & Kolb, J. F. (2012). Dielectric evolution of mammalian cell membranes after exposure to pulsed electric fields. IEEE Transactions on Dielectrics and Electrical Insulation, 19(2), 609–622. DOI: 10.1109/TDEI.2012.6180256

19. Denzi, A., Merla, C., Camilleri, P., Paffi, A., d’Inzeo, G., Apollonio, F., & Liberti, M. (2013). Microdosimetric study for nanosecond pulsed electric fields on a cell circuit model with nucleus. The Journal of membrane biology, 246(10), 761–767. https://doi.org/10.1007/s00232-013-9546-7 PMID: 23595823

20. Lu, W., Wu, K., Hu, X., Xie, X., Ning, J., Wang, C., … & Yang, G. (2017). Theoretical analysis of transmembrane potential of cells exposed to nanosecond pulsed electric field. International journal of radiation biology, 93(2), 231–239. https://doi.org/10.1080/09553002.2017.1230244 PMID: 27586355

21. Kotnik, T., & Miklavčič, D. (2000). Theoretical evaluation of the distributed power dissipation in biological cells exposed to electric fields. Bioelectromagnetics: Journal of the Bioelectromagnetics Society, The Society for Physical Regulation in Biology and Medicine, The European Bioelectromagnetics Association, 21(5), 385–394. PMID: 10899774

22. Merla, C., Paffi, A., Apollonio, F., Leveque, P., d’Inzeo, G., & Liberti, M. (2011). Microdosimetry for nanosecond pulsed electric field applications: a parametric study for a single cell. IEEE Transactions on Biomedical Engineering, 58(5), 1294–1302. DOI: 10.1109/TBME.2010.2104150 PMID: 21216699

23. Merla, C., Denzi, A., Paffi, A., Casciola, M., d’Inzeo, G., Apollonio, F., & Liberti, M. (2012). Novel passive element circuits for microdosimetry of nanosecond pulsed electric fields. IEEE Transactions on Biomedical Engineering, 59(8), 2302–2311. DOI: 10.1109/TBME.2012.2203133 PMID: 22692873

24. Joshi, R. P., & Hu, Q. (2011). Case for applying subnanosecond high-intensity, electrical pulses to biological cells. IEEE Transactions on Biomedical Engineering, 58(10), 2860–2866. DOI: 10.1109/TBME.2011.2161478 PMID: 21937300

25. Salimi, E., Thomson, D. J., & Bridges, G. E. (2013). Membrane dielectric dispersion in nanosecond pulsed electroporation of biological cells. IEEE Transactions on Dielectrics and Electrical Insulation, 20(4), 1256–1265. DOI: 10.1109/TDEI.2013.6571442

26. Hibino, M., Itoh, H., & Kinosita Jr, K. (1993). Time courses of cell electroporation as revealed by submicrosecond imaging of transmembrane potential. Biophysical journal, 64(6), 1789–1800. https://doi.org/10.1016/S0006-3495(93)81550-9 PMID: 8369408

27. Pucihar, G., Miklavcic, D., & Kotnik, T. (2009). A time-dependent numerical model of transmembrane voltage inducement and electroporation of irregularly shaped cells. IEEE Transactions on Biomedical Engineering, 56(5), 1491–1501. DOI: 10.1109/TBME.2009.2014244 PMID: 19203876

28. DeBruin, K. A., & Krassowska, W. (1999). Modeling electroporation in a single cell. I.Effects of field strength and rest potential.Biophysical journal, 77(3), 1213–1224. https://doi.org/10.1016/S0006-3495(99)76973-0 PMID: 10465736

29. Neu, J. C., & Krassowska, W. (1999). Asymptotic model of electroporation. Physical review E, 59(3), 3471. https://doi.org/10.1103/PhysRevE.59.3471

30. Vasilkoski, Z., Esser, A. T., Gowrishankar, T. R., & Weaver, J. C. (2006). Membrane electroporation: The absolute rate equation and nanosecond time scale pore creation. Physical review E, 74(2), 021904.https://doi.org/10.1103/PhysRevE.74.021904 PMID: 17025469

31. Yao, C., Mo, D., Li, C., Sun, C., & Mi, Y. (2007). Study of transmembrane potentials of inner and outer membranes induced by pulsed-electric-field model and simulation. IEEE Transactions on Plasma Science, 35(5), 1541–1549. DOI: 10.1109/TPS.2007.905110 PMID: 19162838

32. Krassowska, W., & Filev, P. D. (2007). Modeling electroporation in a single cell. Biophysical journal, 92(2), 404–417. https://doi.org/10.1529/biophysj.106.094235

33. Sukharev, S. I., Chernomordik, L. V., Abidor, I. G., & Chizmadzhev, Y. A. (1982). 466—Effects of UO22+ ions on the properties of bilayer lipid membranes. Bioelectrochemistry and Bioenergetics, 9(2), 133–140. https://doi.org/10.1016/0302-4598(82)80169-4

